# *C. elegans* chromosomes connect to centrosomes by anchoring into the spindle network

**DOI:** 10.1101/060855

**Authors:** Stefanie Redemann, Johannes Baumgart, Norbert Lindow, Sebastian Fürthauer, Ehssan Nazockdast, Andrea Kratz, Steffen Prohaska, Jan Brugués, Michael Shelley, Thomas Müller-Reichert

## Abstract

The mitotic spindle ensures the faithful segregation of chromosomes. To discover the nature of the crucial centrosome-to-chromosome connection during mitosis, we combined the first large-scale serial electron tomography of whole mitotic spindles in early *C. elegans* embryos with live-cell imaging. Using tomography, we reconstructed the positions of all microtubules in 3D, and identified their plus- and minus-ends. We classified them as kinetochore (KMTs), spindle (SMTs), or astral microtubules (AMTs) according to their positions, and quantified distinct properties of each class. While our light microscopy and mutant studies show that microtubules are nucleated from the centrosomes, we find only a few KMTs are directly connected to the centrosomes. Indeed, by quantitatively analysing several models of microtubule growth, we conclude that minus-ends of KMTs have selectively detached and depolymerized from the centrosome. In toto, our results show that the connection between centrosomes and chromosomes is mediated by an anchoring into the entire spindle network and that any direct connections through KMTs are few and likely very transient.

## Introduction

The mitotic spindle is a dynamic microtubule-based apparatus that ensures the segregation of chromosomes during cell division. Its properties are governed by an array of factors, such as polymerases, depolymerases, motor proteins, cross-linkers and other microtubule-associated proteins^1^. Remarkably, despite the high turnover of microtubules throughout mitosis^2^ the spindle maintains its bipolar structure with the chromosomes at its center and two poles that are separated by the plane of cell division. This stereotypical arrangement is widely believed to mediate the forces between the metaphase plate and the poles that separate sister chromatids during mitosis. In this paper we set out to identify the cytoskeletal ultrastructure in *C. elegans* mitotic spindles that underlies this function, and how this ultrastructure is generated, using a combination of large-scale electron tomography, light microscopy and mathematical modelling.

In all spindles, the microtubule cytoskeleton connects to chromosomes via a special class of microtubules called kinetochore microtubules (KMTs). However, how KMTs bind to chromosomes varies substantially between organisms. In mammals, microtubules attach to monocentric kinetochores that are located at specific sites on the chromosome, whereas nematodes like *C. elegans* have holocentric kinetochores^3^, for which microtubule-binding sites are spread along the entire surface of the chromosomes. If the role of KMTs is to mediate forces between chromosomes and spindle poles they need to somehow connect to the centrosomes. Indeed, that such forces exist in *C. elegans* is evidenced by the maintenance of half-spindle lengths throughout mitosis^4^ and in many perturbations experiments. In budding yeast, single continuous KMTs span the full pole-to-chromosome distance^5^. In mammals, kinetochores and centrosomes are connected by bundles of KMTs, called kinetochore fibres (k-fibres)^1^. It is one aim of our study to identify the nature of the KMT-centrosome connection in *C. elegans*, which is so far unknown.

A related question is the site of KMT-nucleation. Both centrosomes and chromosomes have been proposed as sites of KMT origin^6-9^. In the case of centrosomal origin a radial array of microtubules emanates from centrosomes and those that hit kinetochores can bind and become stabilized as KMTs^10^, ^11^. In the case of chromosomal origin, microtubules instead nucleate around chromosomes and only later attach to kinetochores, as observed in *Xenopus* cell-free extracts^12^. In most systems, the origins of KMTs are unclear^13-15^. Furthermore, centrosomal and chromosomal microtubule nucleation need not be mutually exclusive and may function together during spindle assembly^16^, ^17^ Finally, the nucleation of microtubules in the bulk of the spindle has also been reported^18^, ^19^ Here we address the origin of KMTs in *C. elegans* embryos.

Although *C. elegans* spindles have been widely studied^20^, much remains unknown about the nature and role of the KMTs. While light microscopy provides a dynamic picture of the spindle^14^, ^21-23^, it cannot resolve individual microtubules. Electron microscopy overcomes this limitation though, until now, little quantitative data on the fine structure of mitotic spindles has been published. The available data is mostly limited to *S. cerevisiae*^5^, and partial reconstructions of Ptk2 cells^24^ and early *C. elegans* embryos^25^, ^26^

Here we provide the first full 3D reconstructions of *C. elegans* spindles with single-microtubule resolution using electron tomography. These data allow us, for the first time, to assess the precise locations and spatial relations of all microtubules. We combine this ultrastructural analysis with measurements of microtubule dynamics and show that KMTs in *C. elegans* are nucleated around the centrosomes. Strikingly, KMTs rarely span the entire pole-to-chromosome distance, and using mathematical modelling we show that these findings are consistent with a model in which KMT minus-ends selectively detach from the centrosomes and depolymerise. Our findings imply that the KMT-mediated connection between chromosomes and centrosomes in *C. elegans* spindles is surprisingly transient, which predicts that a direct and permanent connection of chromosomes and centrosomes is not a prerequisite for chromosome segregation.

## Results

We quantitatively analysed the organization of mitotic spindles in the single-cell *C. elegans* embryo. Our electron tomographic approach provided a 3D view of mitotic spindles in metaphase and anaphase (Fig. 1a-d; see Supplementary Video 1 for a full 3D reconstruction of the metaphase spindle; Supplementary Figs. 1 and 2). We analysed data per half spindles (see Table 1 for a summary of all data sets). A half spindle contained 8331 microtubules (Median, *n* = 5), without clear visual differences between metaphase and anaphase. We divided the reconstructed microtubules into three groups: kinetochore microtubules (KMTs), spindle microtubules (SMTs) and astral microtubules (AMTs). All microtubules ending in the ribosome-free zone around the chromosomes were considered as KMTs (Supplementary Video 2)^25^. We detected approximately 227 KMTs per half spindle in metaphase (Median, *n* = 6; Fig. 1 e-f; Supplementary Figure 2 and Supplementary Video 2 for a full 3D reconstruction of the KMTs in Metaphase 1) and 180 KMTs per half spindle in anaphase (Median, *n* = 3; Fig. 1g-h). Non-KMTs that had their centre of mass within a cone with an opening angle of 18.4° towards the chromosomes were classified as SMTs. All others were considered AMTs (Fig. 1i, see also Material and Methods).

**Figure 1.**
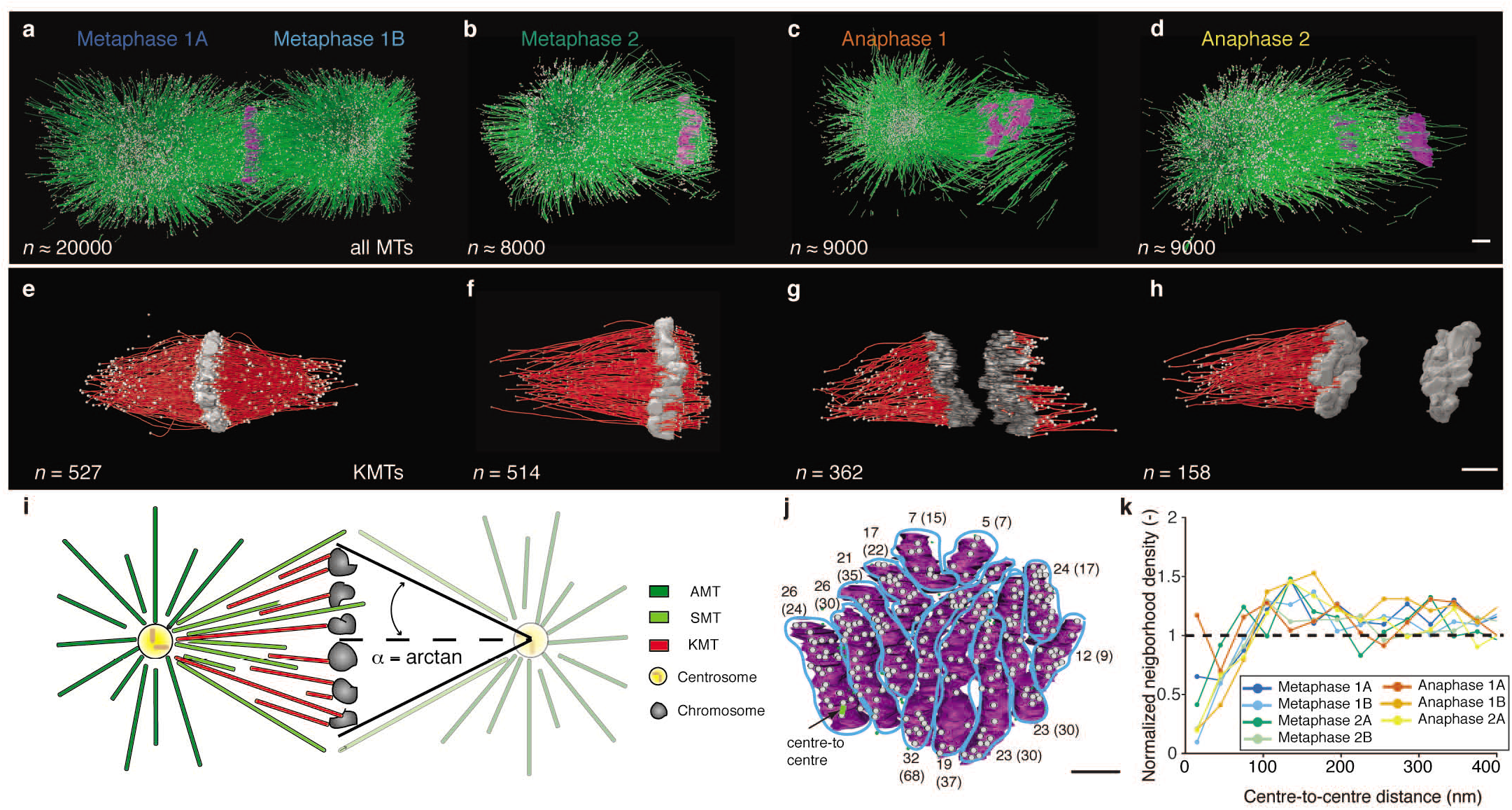
**Three-dimensional reconstruction of spindle and kinetochore microtubules** **a**, Model of microtubules and chromosomes of a full metaphase spindle. **b**, Model of a half spindle in metaphase. **c-d**, Models of half spindles in anaphase. **e-h**, Corresponding 3D models of KMTs in metaphase and anaphase of the reconstructions as shown in **a-d.** The number of microtubules for each reconstruction is indicated. Scale bar, 1 **μ**m. **i**, Schematic diagram illustrating the different microtubule classes (left half) and the geometry of the cone with the indicated opening angle (right half). **j**, End-on view of a metaphase plate 1A. Microtubule attachment to individual chromosomes from each pole is indicated by grey dots. As an example, the green line indicates a centre-to-centre distance between two attachment sites. The numbers of microtubules attaching from the visible pole per chromosome are indicated, the numbers for the opposite poles (metaphase 1B) are given in brackets. Scale bar, 1 **μ**m. **k**, Neighbour density analysis of KMT attachment sites. The radial distribution function is normalized by a random seeding with the same density and geometry. The dashed line indicates the average of random points.

**Figure 2.**
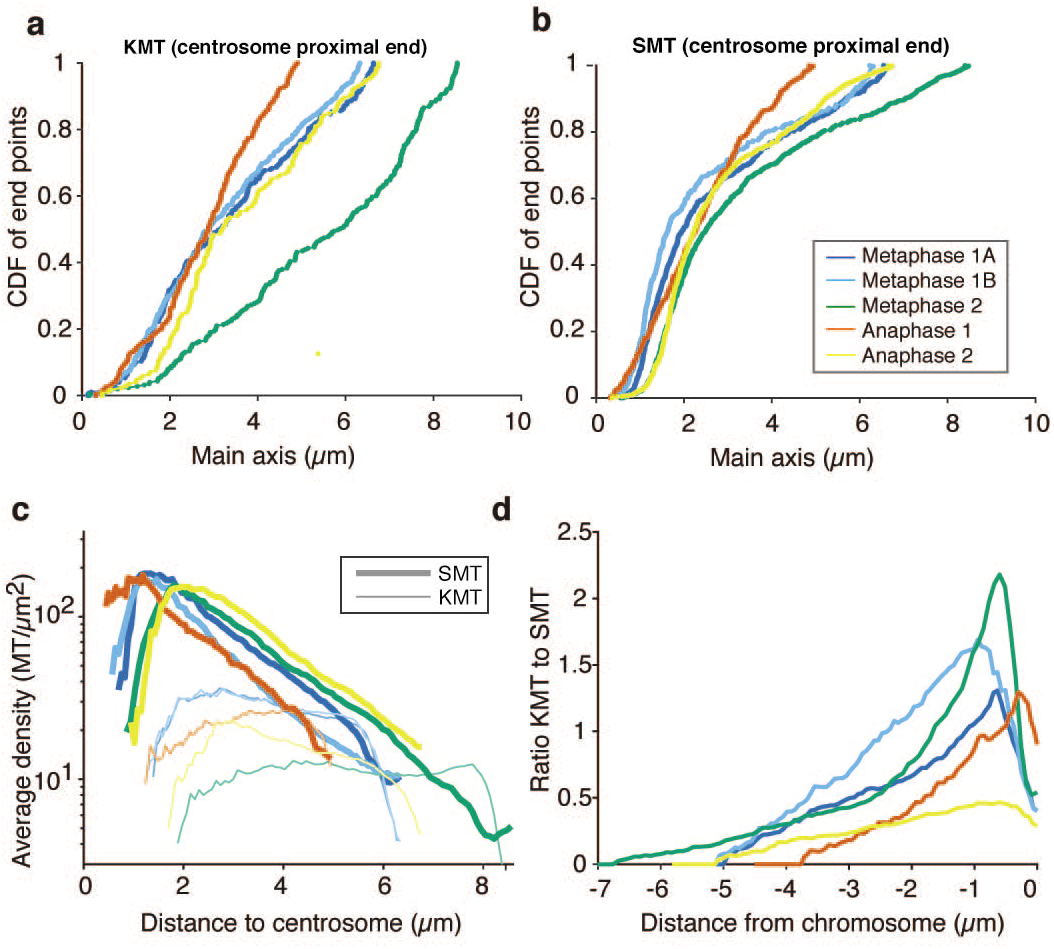
**Analysis of endpoint position and density of kinetochore microtubules** **a**, Plot showing the fraction of ends of SMTs located within a region around the centrosome. **b**, Fraction of ends of KMTs located within a region around the centrosome. **c**, Density of KMTs and SMTs along the half spindle axis from the centrosome to chromosomes measured by counting the microtubules crossing a plane at a certain position on the axis. **d**, Ratio of KMTs to SMTs along the half spindle axis from the centrosome to chromosomes aligned at the chromosomes.

**Table 1.** **Summary of data sets as used throughout this study** This table lists all half spindle data sets and the specific analyses conducted for each data set.

|  | **Meta-phase 1A** | **Meta-phase 1B** | **Meta-phase 2A** | **Meta-phase 2B** | **Meta-phase 3A** | **Meta-phase 3B** | **Ana-phase 1A** | **Ana-phase 1B** | **Ana-phase 2A** |
| --- | --- | --- | --- | --- | --- | --- | --- | --- | --- |
| **Number of AMTs** | 9400 | 9243 | 5713 |  |  |  | 6558 |  | 7356 |
| **Number of SMTs** | 680 | 421 | 727 |  |  |  | 524 |  | 818 |
| **Number of KMTs** | 272 | 310 | 227 | 232 | 152 | 237 | 214 | 181 | 157 |
| **Chrom. attachment** | Figures 1j, S3a, S4a | Figures 1j, S3a, S4b | Figures S3a, S4c | Figures S3a, S4d | Figure S4e | Figure S4f |  |  |  |
| **Neighbor density on chrom.** | Figure 1k | Figure 1k | Figure 1k | Figure 1k |  |  | Figure 1k | Figure 1k | Figure 1k |
| **Density on chrom.** | Figure S3b | Figure S3b | Figure S3b | Figure S3b |  |  | Figure S3b | Figure S3b | Figure S3b |
| **CDF of MT ends** | Figures 2a,b | Figures 2a,b | Figures 2a,b |  |  |  | Figures 2a,b |  | Figures 2a,b |
| **Density/ratio along spindle** | Figures 2c,d | Figures 2c,d | Figures 2c,d |  |  |  | Figures 2c,d |  | Figures 2c,d |
| **Length distribution** | Figures 3a,b,c<br>S11a,b,c | Figures 3a,b,c<br>S11a,b,c | Figures 3a,b,c<br>S11a,b,c |  |  |  | Figures 3a,b,c<br>S11a,b,c |  | Figures 3a,b,c<br>S11a,b,c |
| **Position of short MTs** | Figure 3d | Figure 3d | Figure 3d |  |  |  | Figure 3d |  | Figure 3d |
| **Neighbor density along spindle** | Figures 5c,d | Figures 5c,d | Figures 5c,d |  |  |  | Figures 5c,d |  | Figures 5c,d |
| **Network analysis** | Figures 5e,f | Figures 5e,f | Figures 5e,f |  |  |  | Figures 5e,f |  | Figures 5e,f |
| **open/closed KMT ends** | Figures S6a,b | Figures S6a,b |  |  | Figures S6a,b |  | Figures S6a,b |  | Figures S6a,b |

### Kinetochore microtubules randomly attach to holocentric chromosomes

We first used our data to investigate the distribution of the KMT attachment sites on chromosomes. To this end, we projected the positions of all attached KMT ends on to the plane of cell division. There were 6 to 50 KMTs attaching to each of the twelve chromosomes per pole-facing side (Fig. 1j). Despite the larger kinetochore region, this is surprisingly close to the number of KMTs attaching to the monocentric mammalian kinetochore^27^. We found that the number of attached KMTs correlated with the area of the chromosomes (Supplementary Fig. 3a, Pearson’s correlation coefficient is indicated). The average density of KMTs on the metaphase plates of the half spindles was from 16 to 27 microtubules/**μ**m^2^. Within each dataset the KMT density was nearly constant (Supplementary Fig. 3b). We next asked whether the typical distance between KMT ends on chromosomes was random or followed a pattern that might reveal the existence of preferred attachment sites on the chromosomes. We found the attachment sites to be randomly distributed, with a slight preference towards a spacing of about 127 ± 4 nm (s.e.m., *n* = 7 spindle halves) between two individual KMT ends (Fig. 1k). This weak preferred spacing can arise from the fact that microtubules or their attachment apparati cannot overlap, i.e. they have excluded volume interaction^28^. However, those sites are distributed along the entire length of the chromosomes (Supplementary Fig. 4). We conclude that KMTs in *C. elegans* do not bundle up to form k-fibres. This is consistent with visual inspection of the tomography data.

### Most kinetochore microtubules ends are far from the centrosomes

We next asked whether all KMTs are directly connected to the centrosomes. To answer this question we analysed the distribution of distances of the microtubules’ pole-facing ends from their mother centrioles. In this regard, KMTs are very different from SMTs, as seen in their cumulative distribution functions (CDFs; Fig. 2a-b). The nearly linear CDF for KMTs suggests a nearly uniform distribution of KMT end positions from the centrosome. Conversely, the rapid rise, then levelling, in the SMT CDF shows that SMT ends are mostly clustered near centrosomes. From the CDFs we find that only 22 ± 4 % (s.e.m., *n* = 5 half-spindles) of the KMT ends were located within 2 **μ**m of their corresponding mother centriole (Fig. 2a), while for SMTs, the fraction was 46 ± 4 % (s.e.m., *n* = 5 half-spindles; Fig. 2b). This suggested that the majority of KMTs do not make contact with the centrosomes. In addition, this result prompted us to measure the density of KMTs to SMTs (Fig. 2c) and their ratio along the half spindle axis, which is approximately 6.5 **μ**m in length (Fig. 2d). The ratio of the number density of KMTs to SMTs decreases from chromosomes to poles, dropping from more than one to zero. This further supported the finding that few KMTs span the full distance from chromosomes to centrosomes.

### KMTs have distinctly different length distributions from other microtubules

The centrosome-proximal end position of KMTs and the change in KMT/SMT ratio along the spindle axis suggested a difference in the properties of KMTs versus SMTs and AMTs. In order to address this we analysed the length distribution of the different microtubule classes showing that the three classes of microtubules displayed indeed their own distinct length distributions. AMTs had an exponential length distribution (Fig. 3a). The length distribution of SMTs was exponential for shorter lengths (up to 2 **μ**m), similar to AMTs, followed by a flatter distribution up to about 5-7 **μ**m (Fig. 3b). Very differently, KMTs showed an apparently uniform length distribution, with only a few short microtubules in their population (Fig. 3c; see also Supplementary Fig. 5 for a fit of the length distributions). In summary, this suggests that a different process than those for AMTs and SMTs governs the KMT length distribution. Exponential length distributions as found for AMTs and SMTs are typical of dynamic instability kinetics^29-31^. A uniform length distribution of KMTs, however, indicates a difference in dynamics and possibly a higher stability of the plus-ends against catastrophe.

**Figure 3.**
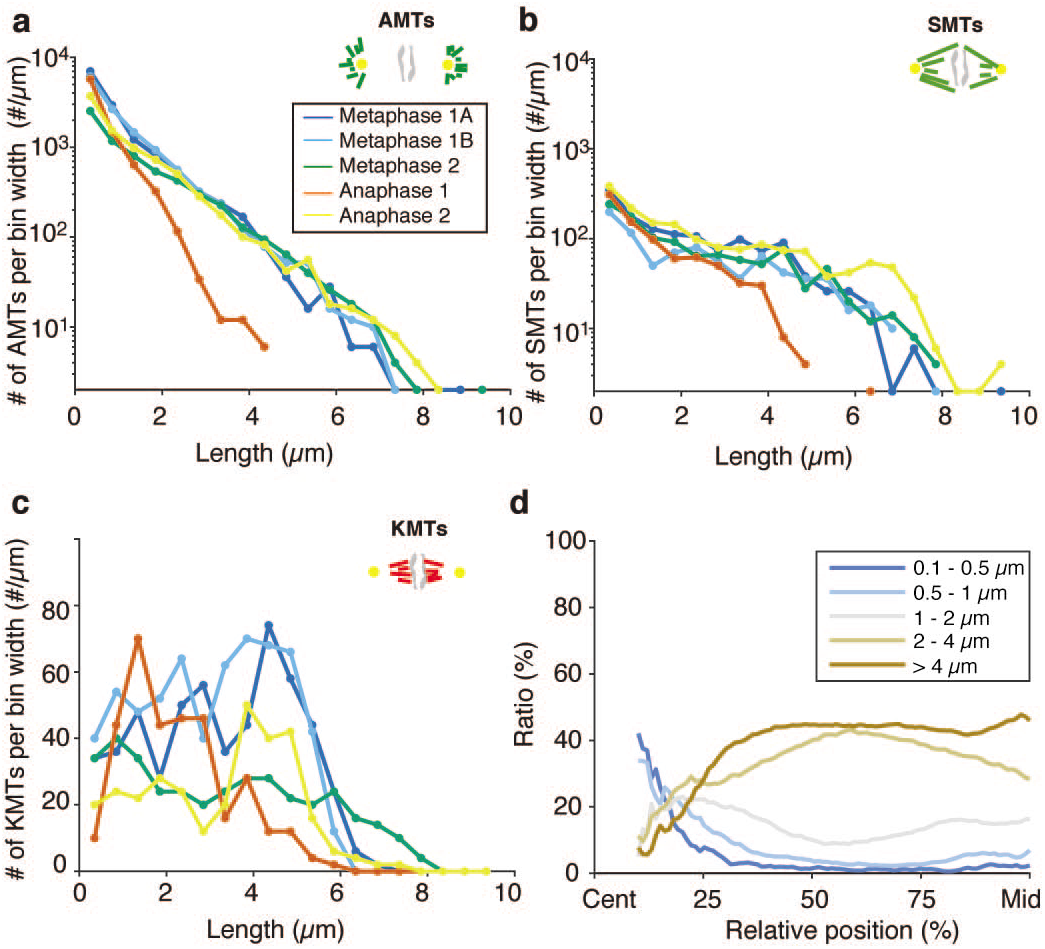
**Microtubule length distributions** **a**, Length distribution of AMTs. **b**, Length distribution of SMTs. **c**, Length distribution of KMTs. **d**, Fraction of SMTs and AMTs within distinct length groups (as indicated by colours, average over all data sets) to all microtubules along the spindle axis from centrosomes to chromosomes.

### Kinetochore microtubules are nucleated at centrosomes

The centrosome in the *C. elegans* mitotic embryo is a major site of microtubule nucleation. However, the KMTs in our reconstructions were not directly connected to centrosomes. This raised the question about the origin of KMTs. To investigate this, we looked at the end-morphologies of KMTs, as an indication for their dynamic state^32-35^. In our reconstructions we distinguished open and closed ends of KMTs (Supplementary Fig. 6a), however, about 40% of the KMT ends could not be unambiguously classified. Analysing the annotated ends, we found that about 71% (n = 766) of those KMT ends at chromosomes in metaphase and 79% (n = 340) of KMT ends in anaphase displayed an open-end conformation with flared ends (Supplementary Fig. 6b). This is consistent with earlier findings^25^, ^26^ Furthermore, 38 % (n = 725) of the pole-facing ends of KMTs in metaphase and 41% (n = 340) of KMT ends in anaphase were open. Analysing only those KMTs with both end morphologies clearly identified, we found that the majority of such KMTs had two open ends (Supplementary Fig. 6c). Since open ends are thought to indicate either growth or shrinkage, our data suggest that most of the KMTs have two dynamic ends. In contrast, closed ends most likely indicate the minus end of a microtubule.

We then analysed the position of microtubules according to their length (Fig. 3d). We found that the majority of short SMTs (below 1 **μ**m) in metaphase and anaphase were located near the centrosomes. This suggests that most nucleation happens near the centrosomes. However, short KMTs are only found near chromosomes, but are not especially prevalent in that population. We thus asked whether KMTs, unlike the majority of microtubules, nucleate at chromosomes. To investigate this, we analysed the formation of KMTs around chromosomes in one-cell embryos in prometaphase (Supplementary Fig. 7a) and two-cell *zyg-1(RNAi)* embryos with monopolar spindles (Supplementary Fig. 7b-d)^36^. Firstly, in both conditions we could not detect any short microtubules on or around chromosomes. Secondly, if microtubules were nucleating around chromatin in the two-cell *zyg-1(RNAi)* embryo, one might expect to see KMTs at the outer side of the metaphase plate (i.e. the side not connected to the spindle pole), which we do not. Hence, we conclude that the chromosomes are not the site of KMT nucleation.

### Microtubules grow unidirectionally away from centrosomes and show different dynamics inside the spindle

As the polarity of individual microtubules cannot be clearly determined in our tomograms, we turned to light microscopy to infer the direction of microtubule growth within the spindle. We visualized the motion of growing microtubule plus-ends by live-cell imaging of EBP-2, which specifically binds to the polymerising microtubule plus-ends (Fig. 4)^37^. The mitotic spindle is a crowded environment preventing the tracking of individual EBP-2 comets. Therefore, we developed a novel method to analyse the EBP-2 velocity within the spindle based on spatial-temporal correlation (see also Material and Methods). We analysed four different regions: within the spindle, at chromosomes, and within a central (inner astral) and a peripheral (outer astral) region of the centrosome (Fig. 4a, Supplementary Video 3). The estimated velocity of the comets was 0.34 ± 0.02 **μ**m/s (s.e.m., *n* = 8 half spindles) in the spindle, 0.49 ± 0.04 **μ**m/s (s.e.m., *n* = 8 half spindles) at chromosomes, and 0.27 ± 0.03 **μ**m/s (s.e.m., *n* = 8 half spindles) in the central region around the centrosome (Fig. 4b). In contrast, we estimated a velocity of about 0.73 ± 0.02 **μ**m/s (s.e.m., *n* = 8 half spindles) in the periphery of the centrosome, suggesting different microtubule dynamics inside spindles than outside of spindles. Additionally we analysed the direction of EBP-2 comets. This showed that most comets move away from the centrosomes and towards the chromosomes (Fig. 4b), indicating that the majority of minus-ends of microtubules in *C. elegans* spindles are located at the centrosomes, whereas plus-ends grow towards the chromosomes. We challenged this finding by performing laser microsurgery to ablate microtubules within the spindle and so measure their polarity by generating new microtubule plus and minus ends^23^. Microsurgery resulted in the formation of a single wave of depolymerisation of the newly created microtubule plus-ends towards the centrosome (Supplementary Video 4). This indicates that microtubules within the spindle have the same polarity, with the minus-ends oriented towards the poles and the plus-ends facing the chromosomes, thus confirming our EBP-2 data. By combining the dynamic data with the ultrastructural data we are able to determine the position of minus-ends as well as plus-ends within the mitotic spindle.

**Figure 4.**
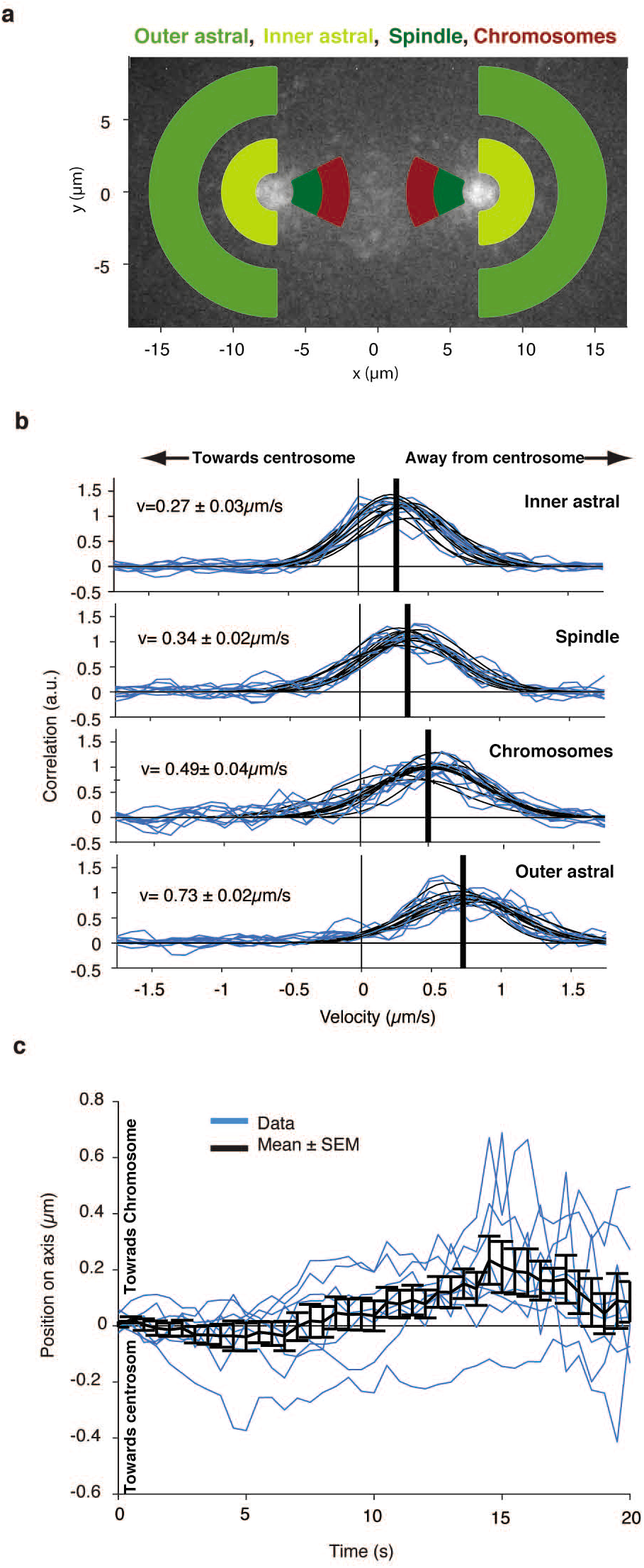
**Directionality and growth velocity of kinetochore microtubules** **a**, Schematic image of different regions used for the analysis of EBP-2. **b**, Cross-correlation of EBP-2 comets for Δt = 0.6 s (blue lines) and measured in the regions as indicated in **a.** The velocity (thick black lines) is estimated by the average over all center positions of the respective Gaussian fits (thin black lines) Velocity and directionality of EBP-2 comets are indicated. **c**, Position of the lowest intensity of the bleach mark over time. Values for the different datasets are shown in blue, the mean (± s.e.m.) is shown in black. Positive is towards the chromosomes.

### Chromosome-bound KMT ends are relatively static

After having established that SMTs grow from their plus ends towards the chromosomes, we sought to understand the behaviour of KMT plus ends. For this, we measured the dynamics of microtubules by FRAP (fluorescence recovery after photo-bleaching) experiments. We bleached a small stripe of approximately 2 **μ**m width in an area close to the chromosomes in metaphase (Supplementary Video 5). To infer the dynamics of the KMT plus ends, which are bound to chromosomes, we measured whether the bleach mark moved (Fig. 4c). Our analysis showed a weak bias of the photo-bleached region for moving towards the chromosomes, although the velocity detected is 0.029 ± 0.005 **μ**m/s and thus close to our detection limit. However, this finding ruled out that KMTs are growing through polymerization at or around chromosomes, since this would result in a motion of the photo-bleached region away from the chromosomes at a velocity that is comparable to the microtubule growth velocity. If anything, the small bias in the opposite direction is consistent with a slow microtubule flux within the *C. elegans* spindle.

### Microtubules in the mitotic spindle are indirectly coupled

Our observation that the majority of KMTs did not reach the centrosome raised the question of how a strong mechanical connection between chromosomes and centrosomes can be achieved during mitosis. Because KMTs may indirectly connect chromosomes to centrosomes, we searched for potential locations of microtubule-microtubule interactions. For such a quantitative network analysis we considered the following parameters: the centre-to-centre distance between two microtubules, the angle between microtubules, and the distance between the pole-proximal end of a non-kinetochore microtubule and the centrosome (Fig. 5a). We started with a neighbour density analysis by measuring the centre-to-centre distance of all microtubules crossing a plane at two distinct positions, at 25% and 75%, along the axis of the half spindle length (Fig. 5b). In comparison to randomly placed microtubules, this analysis revealed an increased frequency of microtubules with a centre-to-centre distance of 55 ± 4 nm at 25% as well as at 75% half spindle length (Fig. 5c-d). This indicates a weak clustering. The measured distances between the microtubules are comparable to the size of microtubule-associated proteins or molecular motors^24^, ^38^, ^39^. However, another possibility is that microtubule-to-microtubule connections are established by cytoplasmic flow and viscous coupling. Moreover the viscous drag forces between nearby microtubules will further couple them mechanically.

**Figure 5.**
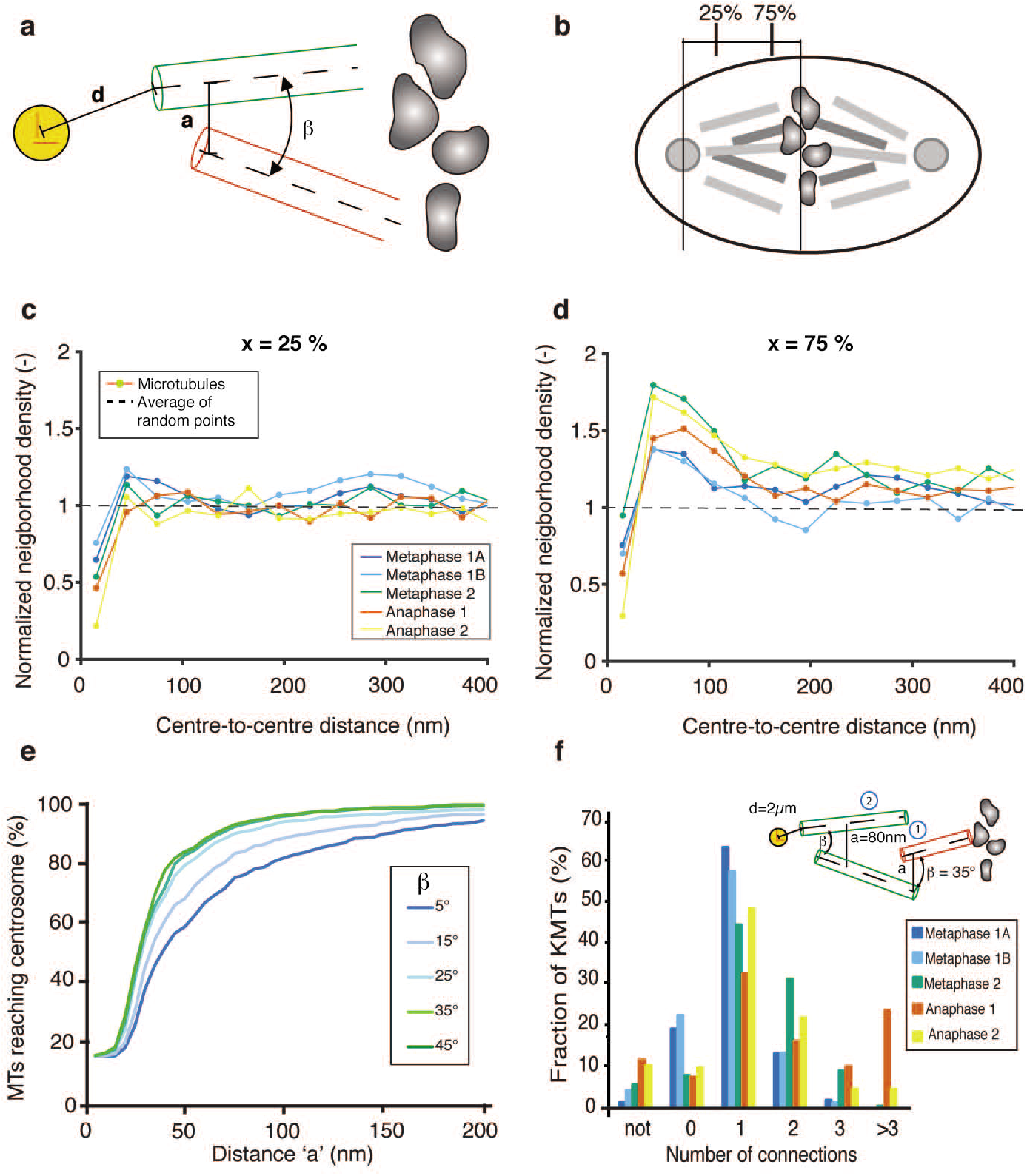
Relative arrangement of kinetochore and spindle microtubules **a**, Parameters for the characterization of microtubule-microtubule interactions. *d:*, distance from centrosome centre to a microtubule (green); *a:* closest centre-to-centre distance between two microtubules (green and red); and β: angle between two microtubules. **b**, Illustration of the positions of 25 % and 75 % of half-spindle length. **c-d**, Neighbourhood density of microtubules at 25 % and 75 % half spindle length for the normalized radial distribution function normalized by random points with the same density on the same geometry. **e**, Percentage of KMTs that can potentially connect to the centrosome as a function of interaction parameters *a* and β. The distance to the centriole *d* is set to 2 **μ**m **f**, Number of interactions necessary to link a KMT to the centrosome for a specific set of parameters (here *a* = 80nm, α = 35°). *‘not’* indicates the number of microtubules that cannot establish a connection, *0’* represents the microtubules that directly connect to the centrosome. A cartoon illustrating a KMT that needs two connections is shown in the inset.

In the light of a possible indirect chromosome-to-centrosome connection we further aimed to analyse the network capabilities of KMTs and SMTs and used the interaction distance and the interaction angle to describe possible microtubule-microtubule interactions. We plotted the fraction of KMTs that are able to connect to the centrosome by multiple interactions. For different interaction angles (5-45°), we plotted the fraction of KMTs reaching the centrosome within a radius of 2 **μ**m as a function of increasing interaction distance (Fig. 5e). This analysis showed that the majority of KMTs could be connected to the centrosome by interacting with SMTs at a 30-50 nm distance, with an interaction angle of 30-45°. By counting the number of interactions that were needed to reach the centrosome, we show that two interactions are typically sufficient to establish a connection to the centrosome in metaphase and anaphase (Fig. 5f). Alternatively, a single KMT might be sufficient for chromosome segregation as shown in budding yeast^5^. Along this line, we found that on average 1-3 KMTs per chromosome in metaphase and 1-2 KMTs in anaphase are directly connected to the centrosome (Fig. 5f). For this analysis we counted KMTs, which had their minus end 2 **μ**m and closer to the mother centriole. However, the outcome of this analysis strongly depends on the set distance of the microtubule ends to the mother centriole. Along this line, within a radius of 1.2**μ**m from the mother centriole on average less than one KMT in metaphase and anaphase directly connect to the centrosome. Both results imply an indirect centrosome-to-chromosome connection and the existence of a spindle network based on KMT and SMT interaction.

### KMT minus-end dynamics is required to maintain observed spindle organization

By combining 3D electron tomography and light microscopy we showed that the KMTs’ length distribution (nearly uniform) is distinct from the SMTs’ (exponential), that the vast majority of microtubules are nucleated near the centrosomes, and that hardly any KMTs span the entire distance from chromosome to centrosome. Moreover, we found that microtubule flux is small. We next sought, by using stochastic simulations of different scenarios of KMT attachment and detachment (Fig. 6), to understand what microtubule dynamics could generate these data.

**Figure 6.**
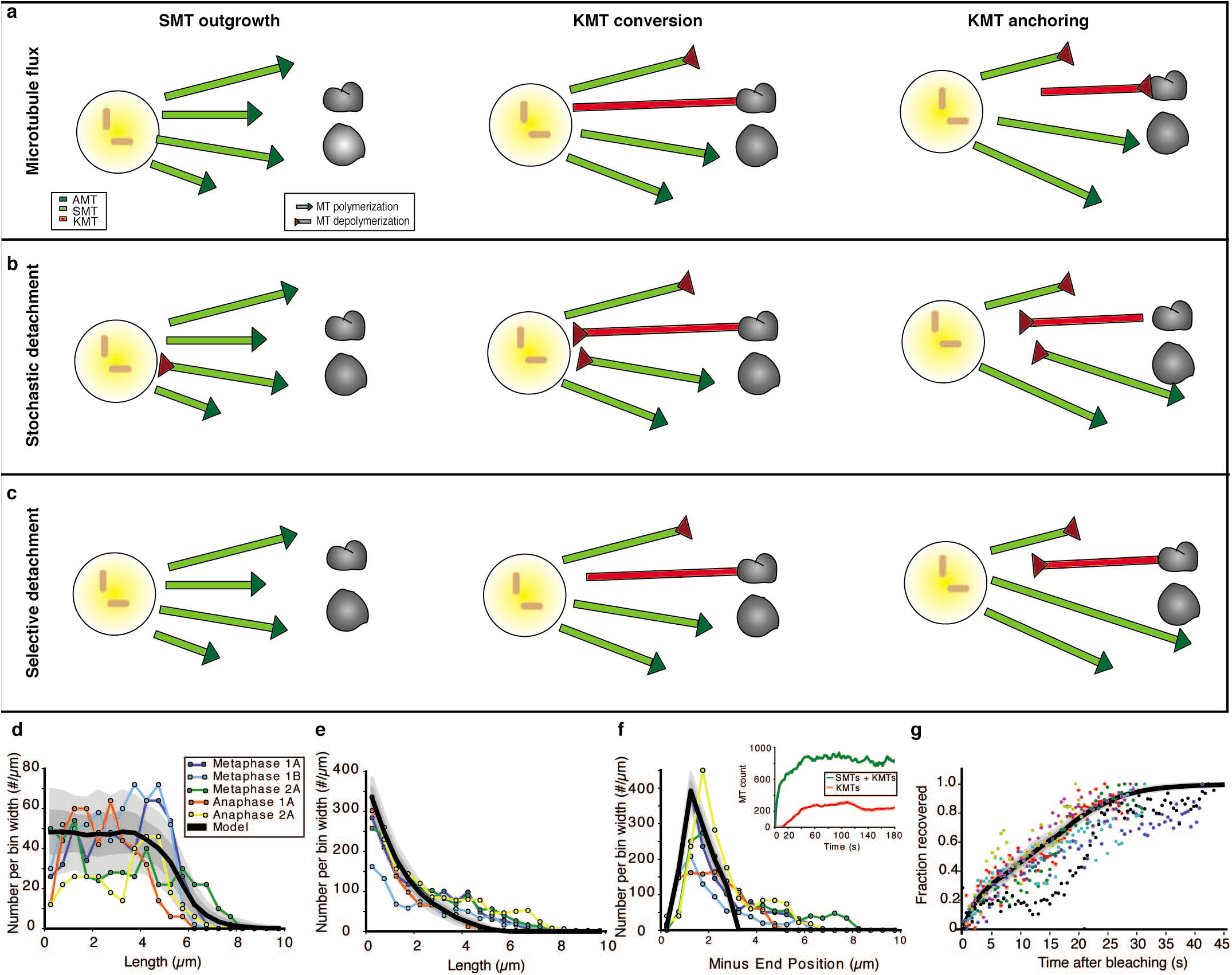
**Models of kinetochore microtubule formation in *C. elegans*** **a**, *Microtubule flux model*: SMTs grow out from the centrosome and will either undergo catastrophe or, upon connecting to a kinetochore, become a KMT. KMTs will then transition to a shrinking state, in which they depolymerise from their plus-end. The plus-end of the KMT remains connected to the kinetochore during depolymerisation. **b**, *Stochastic detachment model*, SMTs grow out from the centrosome and will either undergo catastrophe or, upon connecting to a kinetochore, become a KMT. In this model SMTs as well as KMTs can also depolymerise from their minus-end during their lifetime. **c**, *Selective Detachment Model*, SMTs grow out from the centrosome and will either undergo catastrophe or, upon connecting to a kinetochore become a KMT. KMTs will then transition to a shrinking state, in which they depolymerise from their minus-end. **d**, Results from the selective microtubule detachment model for KMT length distribution. **e**, SMT length distribution. **f**, SMT minus end distribution. The coloured lines show measured EM data. The inset in **f** shows the time-course of the total microtubule number (green) and KMT number (red) for a typical instance of the simulation. **g**, Comparison of experimental FRAP data on microtubule recovery (individual measurement are shown in different colours) with the simulated FRAP data based on the *selective detachment model.* For **d, e, f, g** we display the long time expectation value of the model (solid black line) plus one (dark grey shaded region) and two (light grey shaded region) standard deviations.

In our modeling, we assigned each microtubule a nucleation time from a Poisson process with a nucleation rate *R*, and initial minus-end position within 3**μ**m of the centrosome based on the measured distribution of SMT minus-ends and the position of short microtubules within the spindle (see Fig. 2b and Fig. 3d). The nucleation rate was adjusted such that the steady-state emerging from our simulations had a number of KMTs compatible with our experimental findings (Fig. 1e-h). SMTs grew from their plusends with a velocity *V*_*g*_ = 0.4 *μ*m/s (as measured; see Fig. 4b), until they either underwent catastrophe with rate κ = 0.25 s^-1^, as estimated from the decay of the length distribution of short SMTs (see Material and Methods), or reached the metaphase plate. Note that for simplicity we assumed that catastrophe from a free plus-end immediately destroys a microtubule and so we did not track SMT depolymerization explicitly. SMTs that reached the chromosomes, which were positioned *L* = 6.5 *μ*m away from the centrosomes (as measured from ultrastructure), attached and became KMTs and could no longer undergo catastrophe from their plus-ends. Finally, KMTs only rarely spanned the entire centrosome-to-chromosome distance (Figs. 2a, 3c, 5f), which suggested that upon becoming KMTs microtubules rapidly switched to a depolymerizing state.

Within these constraints we formulated three models of KMT and SMT dynamics, which we called *flux*, *stochastic detachment*, and *selective detachment models*, respectively (see Fig. 6 and model flowcharts in Supplementary Fig. 8, see Table 2 for parameters). In the *flux model*, microtubule plus-ends switched deterministically to shrinking upon becoming KMTs, while staying stably attached to the chromosomes (see Fig. 6a), and the minus-end became detached from the centrosome. We took the overall plus-end shrinking velocity *V*_*d*_ = 0.03 *μ*m/s in accordance with our FRAP measurements (Fig. 4c). In this model there were no adjustable parameters. To compare the model to the experimental data, we ran the simulation sufficiently long to reach statistical steady state, which was then sampled several times, over long times, to obtain an expectation and standard deviation for the extracted distributions.

**Table 2.** **Parameters for the three stochastic models** The adjustable parameters of the simulations are set in bold italic. All other values in the table were estimated from experimental observations.

|  | Flux model | Stochastic detachment model | Selective detachment model |
| --- | --- | --- | --- |
| Growth Velocity $v_g$ | 0.4 $\mu\text{m/s}$ | 0.4 $\mu\text{m/s}$ | 0.4 $\mu\text{m/s}$ |
| Depolymerization Velocity $v_d$ | 0.02 $\mu\text{m/s}$ | ***0.45 $\mu\text{m/s}$*** | ***0.17 $\mu\text{m/s}$*** |
| Centrosome-Chromosome Distance $L$ | 6.5 $\mu\text{m}$ | 6.5 $\mu\text{m}$ | 6.5 $\mu\text{m}$ |
| Catastrophe rate | 0.25 Hz | 0.25 Hz | 0.25 Hz |
| Switching rate $r$ for KMTs | Instantaneous | ***0.2 Hz*** | 0.5 Hz |
| Switching rate $r$ for SMTs | 0 | ***0.2 Hz*** | 0 |

The *flux model* produced a KMT length distribution consistent with the data (Supplementary Fig. 9a), but given the low shrinking velocity *V*_*d*_ and the constraint of producing the observed number of KMTs it underestimated the number of SMTs by a factor of five (relative to observation, and as reflected in length and minus-end position frequencies plotted in Supplementary Fig. 9b-c). Furthermore, in the *flux model* a *de-novo* generated spindle took more than 5 minutes to reach its steady state, which is long compared to the typical duration of metaphase in *C. elegans* (Supplementary Fig. 9c, inset). We concluded that microtubule plus-end shrinking alone is insufficient to explain the data. This suggested that microtubule minus-ends in the spindle are dynamic.

We now investigated models where KMTs shrink from their minus-ends, and not their plus-ends. In the *stochastic detachment model* (Fig. 6b) all microtubule minusends, whether SMT or KMT, switched stochastically, with rate *r*, to a shrinking state, and moved away from the centrosome. In the *selective detachment model* (Fig. 6c) only KMT minus-ends could switch, while SMT minus-ends remained unconditionally stable. Furthermore, in the *stochastic detachment model* KMT plus-ends kept growing against the chromosomes even after the minus end started to shrink, while in the *selective detachment model* plus ends attached to chromosomes stopped growing after the onset of minus-end depolymerization. In the detachment models the minus-end depolymerization velocity *V*_*d*_ and the switching rate *r* were adjustable parameters. As with the *flux model*, the simulations were evolved to statistical steady-state, after which the desired distributions were extracted.

We found that both models could be tuned to produce numbers of both KMTs and SMTs close to experiments (Fig. 6d-e, Supplementary Fig. 10a-c), while reaching steady-state in under a minute, which is compatible with the duration of mitosis in *C. elegans* (Fig. 6f, inset; Supplementary Fig. 10c, inset). However, the *selective detachment* model captured far better the shapes of the KMT length (Fig. 6d, Supplementary Fig. 10a) and the distribution of SMT minus-end position distributions (Fig. 6f; Supplementary Fig. 10c).

We next asked whether the models that we inferred from static tomographic data would also account for spindle dynamics. For this, we used our models to predict the FRAP dynamics of a box of photobleached spindle material with a width of 1 *μ*m at a distance of 2.5*μ*m from the chromosomes. We then plotted the predictions of our models and compared them to the mean intensity measurements of our FRAP experiment (Fig. 6g, Supplementary Fig. 9d, 10d). The recovery rate in metaphase as measured by FRAP was approximately t_1/2_ = 21.4 (19.7, 23.2) s (95 % CI, *n* = 7 spindles), in agreement with previously reported data^40^. We find that the *selective detachment* model quantitatively captures our FRAP data, whereas the *stochastic detachment* and *flux* models do not. For the *selective detachment model*, the recovery curve is the sum of a fast (5s) exponential contribution from recovering SMTs and a slower (20s) linear contribution from recovering KMTs.

Our findings imply that KMTs are transient, despite their plus ends being stabilized against catastrophe. This implies that the spindle can recover its unperturbed structure rapidly (i.e. within 20s) even after drastic disruptions, such as local laser ablation. Our modelling further suggests that selective destabilization of KMT minus-ends is required for the observed spindle structure. We predict that an experiment inhibiting minus-end depolymerization would measure a KMT length distribution that was clustered around the centrosome-to-chromosome distance, instead of uniform, and observe the number of KMTs increase linearly as time progresses from metaphase to anaphase (assuming a wealth of KMT binding sites). In contrast, for an experiment with all microtubule minus-ends instead rendered unstable, we predict a KMT length distribution that is exponentially decaying rather than uniform.

## Discussion

The mitotic spindle ensures the faithful segregation of chromosomes, which requires that a connection between centrosomes and chromosomes be established and maintained throughout mitosis. Prior to our work, it was largely unknown how the ultrastructure of the microtubule cytoskeleton supports this role and provides a coupling which resist the forces acting on the spindle during mitosis^41^, yet robust against even drastic perturbations^42^. To address this, we provided the first complete ultrastructures of five *C. elegans* mitotic spindle halves, which together with dynamic light microscopy revealed the origin of KMTs, the nature of the connection between chromosomes and centrosomes, and enabled us to formulate a mathematical model for the establishment and maintenance of spindle architecture.

From electron microscopy and from tracking the dynamics of growing microtubule plus-ends we found that the large majority of microtubules in the *C. elegans* mitotic spindle are nucleated in a small region around the centrosomes. This is strikingly different from spindles in acentrosomal *C. elegans* oocytes where microtubules nucleate around chromosomes^43^, ^44^ It seems that the presence of centrosomes inhibits or outcompetes other pathways of microtubule nucleation at this stage. Indeed, in mutant studies we could not detect microtubules nucleating around the chromosomes. Thus, we conclude that spindle microtubules, including KMTs are nucleated around the centrosomes.

Given the centrosomal origin of KMTs it is striking that in electron microscopy the majority of their minus-ends are remote from the spindle pole. Only 22% of KMTs reach within a distance of 2 *μ*m from the centrioles. If indeed the role of KMTs is to connect centrosomes to chromosomes, this suggests that they do so by anchoring into the spindle network rather than by a direct linkage. In our network analysis we found that most KMTs could connect to the centrosomes by one or two intermediate microtubules, given reasonable assumptions on the size of potential linker molecules. Visualizing these linkers is, however, far beyond our resolution limit. For now, we speculate that the anchoring of KMTs into the spindle network might be supported by mechanisms similar to the ones found to link k-fibres into the spindle network in mammalian cells^45^, ^46^, where dynein seems to be the main crosslinking agent. An indirect centrosome-to-chromosome connection could be further supported by a viscous coupling, as the microtubules within the cytoplasm might be close enough to generate an enhanced viscous drag^47^. Anchoring of KMTs into the spindle network might also explain similarly loose KMT architectures in other organisms, such as crane flies^48^ or the algae *Oedonium*^49^. It will be important to explore the differences between the ‘anchoring’ mechanism we propose here and direct connections, such as the ones observed for instance in Ptk2 cells^24^, and their implications for cell division. We speculate that anchoring into a spindle network can provide stability while the KMTs turnover every 20 seconds. This might be particularly important for spindles that operate under strong external forces, such as the *C. elegans* spindle, which during normal cell division experiences strong pulling forces from the cell cortex, yet maintains its size and shape^42^.

To understand how the anchoring architecture of *C. elegans* mitotic spindles is maintained we turned to mathematical modeling. We found that the detachment of KMTs from the spindle pole in *C. elegans* is most likely explained by selective destabilization of their minus-ends once the plus-ends bind to kinetochores. This detachment could be achieved either mechanically through compressive loads building up on growing KMTs spanning chromosomes and centrosomes, or biochemically by specifically targeting the KMT minus-ends. In any case, our model provides robust predictions of how spindle structure would change in experiments targeting the detachment mechanism. Surprisingly perhaps, our mathematical model also predicts that the lifetime of KMTs is, like SMTs, short relative to the time-scale of mitosis. We speculate that the rapid turnover of all microtubules might be required to maintain a robust yet flexible enough spindle architecture to correct against perturbations, since it allows the spindle to recover from perturbations within about 20 seconds.

Our finding that the connection between centrosomes and chromosomes is supported by anchoring into the spindle network rather than by direct links, together with the observation that the centrosome-to-chromosome distance remains constant throughout anaphase^50^, raises the question how the segregation of the sister chromatids is achieved. It is tempting to speculate that microtubules organized between the segregating chromatids may play an important role during mitotic chromosome segregation, similar to meiotic divisions in *C. elegans* oocytes^43^. This view on the role of inter-chromosomal microtubules is supported by the observation that chromatids in *C. elegans* mitosis can segregate without centrosomes in a CLASP-dependent manner^51^. We strongly believe that a detailed ultrastructural analysis of such inter-chromosomal microtubules is urgently needed to support any further robust discussion on chromosome segregation.

## Materials and Methods

### Worm strains, gene silencing by RNA interference and feeding clones

*C. elegans* strains were cultured as described^52^. All strains were maintained at either 16 °C or 25 °C. The following strains were used in this study: wildtype N2 Bristol; MAs37 (unc-119(ed3) III; [pie-1::epb-2-gfp;unc-119(+)]^53^. RNAi experiments were performed by feeding as described^54^. Worms for *zyg-1* (RNAi) were grown for 24 h at 25 °C on feeding plates. The feeding clone for *zyg-1* (F59E12.2) was provided by A. Hyman (Dresden, Germany).

### Sample preparation for electron microscopy

Wild-type N2 *C. elegans* hermaphrodites were dissected in M9 buffer and single embryos early in mitosis were selected and transferred to cellulose capillary tubes (Leica Microsystems, Vienna, Austria) with an inner diameter of 200 *μ*m. The embryos were observed with a stereomicroscope until either metaphase or anaphase and then immediately cryo-immobilized using an EM PACT2 high-pressure freezer equipped with a rapid transfer system (Leica Microsystems, Vienna, Austria) as previously described^55^. Freeze substitution was performed over 3 d at −90 °C in anhydrous acetone containing 1 % OsO_4_ and 0.1 % uranyl acetate using an automatic freeze substitution machine (EM AFS, Leica Microsystems, Vienna, Austria). Epon/Araldite infiltrated samples were flat embedded in a thin layer of resin, polymerised for 3 d at 60 °C, and selected by light microscopy for re-mounting on dummy blocks. Serial semi-thick sections (300 nm) were cut using an Ultracut UCT Microtome (Leica Microsystems, Vienna, Austria). Sections were collected on Formvar-coated copper slot grids and poststained with 2 % uranyl acetate in 70 % methanol followed by Reynold’s lead citrate^56^.

### Data acquisition by electron tomography

Dual-axis electron tomography was performed as described^57^. Briefly, 15 nm colloidal gold particles (Sigma-Aldrich) were attached to both sides of semi-thick sections collected on copper slot grids to serve as fiducial markers for subsequent image alignment. For electron tomography, series of tilted views were recorded using a TECNAI F30 transmission electron microscope (FEI Company, Eindhoven, The Netherlands) operated at 300 kV. Images were captured every 1.0° over a ±60° range and a pixel size of 2.3 nm using a Gatan US1000 CCD camera (2k × 2k). For each serial section two montages of 2 × 3 frames were collected and combined to a supermontage using the IMOD software package to cover the pole-to-pole distance of the spindles^58^. For image processing the tilted views were aligned using the positions of the colloidal gold particles as fiducial markers. Tomograms were computed for each tilt axis using the R-weighted back-projection algorithm^59^. For double-tilt data sets two montages, each consisting of six tomograms, were aligned to each other and combined to a supermontage^57^. In order to cover a large volume of the pole-to-pole region of each mitotic spindle, we recorded on average 24 consecutive serial sections per spindle.

### Three-dimensional reconstruction and automatic segmentation of microtubules

We used the IMOD software package (http://bio3d.colourado.edu/imod), which contains all of the programs needed for calculating electron tomograms^58^. Reconstructed tomograms were flattened and the two acquired montages of each section were combined to a supermontage using the *edgepatches, fitpatches* and *tomostitch* commands contained in the IMOD package. We applied the Amira software package with an extension to the filament editor of the Amira visualization and data analysis software for the segmentation and automatic tracing of microtubules^60^. We also used the Amira software to stitch the obtained 3D models in *z* to create full volumes of the recorded spindles^61^. The automatic segmentation of the spindle microtubules was followed by a visual inspection of the traced microtubules within the tomograms and correction of the individual microtubule tracings. Corrections included: manual tracing of undetected microtubules, connection of microtubules and deletions of tracing artifacts (e.g. membranes of vesicles). Approximately 5 % of microtubules needed to be corrected.

### Data analysis

Data analysis was performed using either the Amira software (Visualization Sciences Group, Bordeaux, France) or by Matlab (R2015b, The MathWorks Inc., Nitick, USA).

#### A. Neighbourhood density of microtubules

The microtubule neighbourhood densities for 2D slices in comparison to random samples and displacements were computed in two steps. First, uniformly distributed slices were defined along the centrosome-to-chromosomes axis for each half spindle. Additionally, a cone was defined along the same axis, starting at the centre of the mother centriole and opening with an angle of 18.4° towards the chromosomes (Figure 1 i). The intersection area of this cone with each slice thus determined the regions for the microtubule density measurements. Second, the radial distribution function was estimated. For each microtubule point, the local density in a range of radial distances was computed. The mean over all microtubules provided an estimate for the radial distribution function as a neighbourhood density. For the normalization we used 10,000 sets of randomly placed microtubules with the same total number.

#### B. KMT attachment to chromosomes

In order to correlate the number of KMTs attaching to the chromosome surface we assumed the shape of the chromosome surface available for KMT attachment to be a rectangle. This area of each rectangle corresponding to a chromosome was then correlated to the number of KMT attaching to the individual chromosome.

#### C. Length distribution of microtubules

For the analysis of the microtubules length distributions (Fig. 3a-b), we checked whether the microtubules that leave the reconstructed tomographic volume affect our results (approximately 11 *μ*m × 16.5 *μ*m × 6 *μ*m for each half spindle). We removed microtubules with one end point less than 250 nm apart from the boundary of the volume. These microtubules potentially leave the tomographic volume. This had only consequences for the length distribution of the AMTs in terms of the total number and changed only slightly the shape of the distribution (Supplementary Fig. 11). Furthermore, in all analyses, microtubules shorter than 100 nm were excluded to reduce errors due to the minimal tracing length. In addition, the end point type could not always be identified during inspection. The number of unclear end points lies in the range of 2 % and is uniformly distributed over the kinetochore region. Therefore, we do not expect a relevant error in this analysis.

#### D. Network analysis

For the detection of possible interactions in 3D, a three-step algorithm was implemented in Amira. First, for each microtubule, the distance to the centriole was computed and all microtubules with a distance smaller than this distance were marked as connected to the centrosome. It is important to note here that each microtubule is represented as a piece-wise linear curve. For each line segment of a microtubule the distance to the centriole, which is represented as a point, was computed analytically. The distance of a microtubule was defined as the minimum of all segment distances. Second, for each pair of microtubules the distance and the angle were computed. The distance between two microtubules was defined as the minimum of the distances between all their line segments. A 3D grid data structure was used to accelerate these computations. To reduce errors due to local distortions of the microtubules, the angle is defined by the angle between the lines through the start and end points of the microtubules. Third, based on these data an abstract graph was constructed, where each microtubule is represented as a vertex and each interaction (based on thresholds for interaction distance and angle) as an edge. Finally, for each KMT the shortest path to a microtubule marked as connected to the centrosome was computed in the graph using Dijkstra’s algorithm.

### Error analysis of microtubule segmentation and stitching

For the complete imaging, reconstruction, and microtubule segmentation pipeline of a spindle, the following errors needed to be investigated. First, during the data preparation and the imaging process, the tomograms are locally distorted. Furthermore, the exposure of the electron beam causes a shrinking of the sample. During the reconstruction of the microtubules, however, most errors occur in the tracing and matching process. Additionally the data is again distorted in all directions to align the tomograms. We assumed that this distortion primarily compensates the distortion of the imaging process. For the tracing, the error was previously analysed for reconstructions of *C. elegans* centrosomes^62^. Although the spindle data is larger, the tomogram content is similar to the centrosome data sets, and thus we assumed that the error lies in the same range of 5-10%. In addition, the traced microtubules were manually verified. It is more difficult to estimate the error of the matching algorithm^61^, since it depends on the local density and properties of the microtubules. For this reason, the stitched microtubules were manually verified and corrected for all KMTs. The quality of the analysis of the KMTs, therefore, should be influenced only by minor spatial distortions. In order to estimate the overall quality of the stitching, the distribution of microtubules endpoints in z-direction (i.e. normal to the plane of the slice) was analysed by binning the endpoints in z-direction (Supplementary Fig. 12). Bins were fixed to be either inside a section (50 % of slice thickness in z-direction, centred) or across a boundary between sections (25 % of slice thickness in z-direction of either adjacent section). In order to account for a varying section thickness a microtubules endpoint density (in z-direction) was defined by normalizing over the width of these bins. We assume that high quality stitching would result in a smooth curve. However we did detect some peaks within the histograms. Generally most of these peaks are found within the sections. This can be explained by the fact that the boundary regions of a tomogram are often blurry and microtubules are possibly not traced within this area. This would explain systematically lower endpoint number in boundary regions and the saw tooth features in the histograms. This may be especially relevant in regions were microtubules run parallel to boundaries.

### Light microscopy

Worms were dissected in M9 buffer on a coverslip to obtain embryos. The embryos were then transferred to a glass slide with a 2 % agarose pad.

#### A. EBP-2 analysis

Imaging of the EBP-2::GFP comets was carried out on a Nikon TiE spinning disc confocal microscope using a Nikon Plan-Apochromat 60x water-immersion objective and an iXon EM + DU-897 BV back illuminated EMCCD camera (Andor, Belfast, UK). A single plane was acquired every 250 ms with an exposure of 200 ms starting from metaphase until embryos reached telophase using the IQ3 software (Andor, Belfast, UK). We analysed the local velocities of growing microtubule tips labeled by EBP-2. To obtain a robust estimate in the highly crowded spindle, EBP-2 comets were segmented in each frame using the mosaic suite in Fiji^63^. We then analysed the spatial-temporal correlations of the segmented EBP-2 comets along the radial direction. This approach avoids the problem of linking the right EBP-2 mark in subsequent frames in a crowded environment. The initial segmentation is necessary as otherwise the signal-to-noise ratio is not sufficient. The spatial-temporal correlations were computed by first resynthesizing movies, where each identified EBP-2 spot was convolved with a Mexican-hat wavelet. Along the radial direction the size was set to a half pixel size and in the orthogonal direction enlarged by a factor of 4. This ensures that motions along the circumferential direction are still permissible. For the time lag of the spatial-temporal correlations we used 0.6 s and we averaged over all circumferential positions and over the duration of metaphase.

#### B. FRAP analysis

For FRAP experiments we used a Nikon microscope (Yokogawa CSU-X1 Spinning disk; equipped with a 60x 1.2 NA objective, Chamaleon 2-photon laser for ablation and an Andor Ixon Ultra 897 camera). For data analysis the position of the two centrosomes was identified and an intensity profile extracted along this axis. We averaged in the perpendicular direction over a distance of 2 *μ*m. The profiles were aligned along the axis by fitting a Gaussian profile to the intensity peak of chromatin, which was labelled by histone::GFP. The photo-bleached region was fitted by a 2^nd^ order polynomial and the location estimated from the position of the minimum. We used the distance between these two to estimate the velocity of the photo-bleached region with respect to the chromosomes. For the recovery we analysed the amplitude at the centre of the photo-bleached region with respect to the intensity at the mirrored position on the axis.

### Stochastic simulations of KMT formation

We performed stochastic simulations for three different models of microtubule dynamics, which we call the *flux model*, the *stochastic detachment model*, and the *selective detachment model*, respectively (Supplementary Fig. 8). The models were implemented using a standard Gillespie algorithm.

In the following we lay out how the parameters for our stochastic models have been chosen. We first need to specify where and when new microtubules nucleate. Under the assumption that most microtubules are SMTs and that the minus ends of SMTs are mostly immobile, the measured distribution of SMT minus-end positions provides a good estimate for the nucleation positions of microtubules. We thus use this measured distribution to determine the nucleation position of new microtubules in our model. Note that we truncate the measured distribution at a distance of 3 *μ*m from the centrosomes, since minus ends further away are most probably caused by effects that our modeling does not capture. The resulting nucleation profile is shown in Figure 6f.

Before attaching to chromosomes, SMTs only grow at the velocity *V*_*g*_ and catastrophe at the rate *κ*. Thus the length distribution *ψ*(ℓ) of SMT length obeys ∂_*t*_*ψ*(ℓ) = − *V*_*g*_∂_ℓ_*ψ* − *κψ*, which is solved at a steady state by *ψ*(ℓ) = *A exp*(− ℓ(*V_g_K*)). Since *V*_*g*_ = 0.4 **μ**m/s is known from direct measurements (see Fig. 4b), we can infer *κ* by fitting to experiments, and obtain *κ* ≃ 0.25 Hz (see Supplementary Fig. 5).

Furthermore we need to specify the distance *L* from chromosomes to centrosomes, which we take at 6.5 **μ**m in accordance to our ultrastructure data. Finally we need to specify the speed of KMT depolymerization *V*_*d*_ at which microtubules shrink and the rate *r* at which microtubules make the switch from growth to shrinking. For the flux model *V*_*d*_ is bounded by the measured flux velocity of 0.02 **μ**m/s, which is the value we prescribe. With this velocity having the switch from SMT to KMT be deterministic (i. e. *r* goes to infinity) yields the best results. For the stochastic and selective detachment models *r* and *V*_*d*_ are a priori not known. We adjust them to yield best agreement between experiments and data. All of these values are summarized in Table 2.

The three models differ in the following aspects: In the *flux model*, upon becoming KMTs, KMT plus-ends switch deterministically to shrinking at a velocity *V*_*d*_ = 0.02 *μ*m/s. In the *depolymerisation model*, both KMTs and SMTs can switch to depolymerising from their minus ends with a velocity *V*_*d*_ at a rate r. In the *detachment model*, only KMTs can switch to depolymerising from their minus-ends with a velocity *V*_*d*_ at a rate r. While the *flux model* has no adjustable parameters, in the depolymerisation and detachment models the rate *r* and the velocity *V*_*d*_ are unknown. Requiring the ratio of SMTs to KMTs to match experiments and mimicking the shape of the experimentally observed KMT length distribution set both rates.

To compare the outputs of our simulations to the experimental data, we run the simulation sufficiently long to reach a steady-state, and then average over a large number of subsequent steady-state configurations, sampled every thirty seconds to obtain an expectation value and standard deviations for the extracted distributions.

We also extracted predictions for the time course of FRAP experiments from each of our models. In these numerical experiments we specify the position and width of the bleached box, and track the positions of all microtubule segments, bleached or unbleached, that are inside this box. We then calculate the fraction *S(t) of* unbleached MTs *t* time units after the bleaching event. This fraction is given by *S(t)=[M(t)-B(t)]/M(t)*, where *M(t)* is the total mass of microtubules inside the box at time t, and *B(t)* is the total mass of bleached material remaining at time t. We compare *S(t)* directly with the normalized fluorescence intensities from our FRAP measurements (Fig. 6g and Supplementary Figs. 9d and 10d).

## Acknowledgments

We thank F. Jülicher, D. Needleman, and Dr. Sándalo Roldán-Vargas for continuous discussions and A. Hyman for the feeding clone for *zyg-1* (F59E12.2) and continued support. The authors are also grateful to Martin Merkel for microtubule segmentation and the electron microscopy facility at MPI-CBG (Dresden) for technical assistance. The Müller-Reichert lab received funding from the Human Frontier Science Program (RGP 0034/2010), the German Research Foundation (DFG grant MU 1423/8-1) and from the Saxonian State Ministry for Science and the Arts (SMWK). J. Baumgart received funding from the European Comission’s 7^th^ Framework Programme grant Systems Biology of Mitosis (FP7_HEALTH-2009-241548/MitoSys). The Brugués lab acknowledges funding from the Human Frontier Science Program (CDA 74/2014), S. Prohaska was funded by the German Research Foundation (DFG grant PR 1226/4-1) and the FEI Visualization Sciences Group. The Shelley lab acknowledges support from the (USA) National Institutes of Health (1R01GM104976-01), the National Science Foundation (DMS-1463962), and the Human Frontiers Science Program for support of S. Fürthauer.

## Author contributions

This work represents a truly collaborative effort. Each author has contributed significantly to the findings and regular group discussions guided the development of the ideas presented here.

## Figure Legends

**Supplementary Figure 1.**
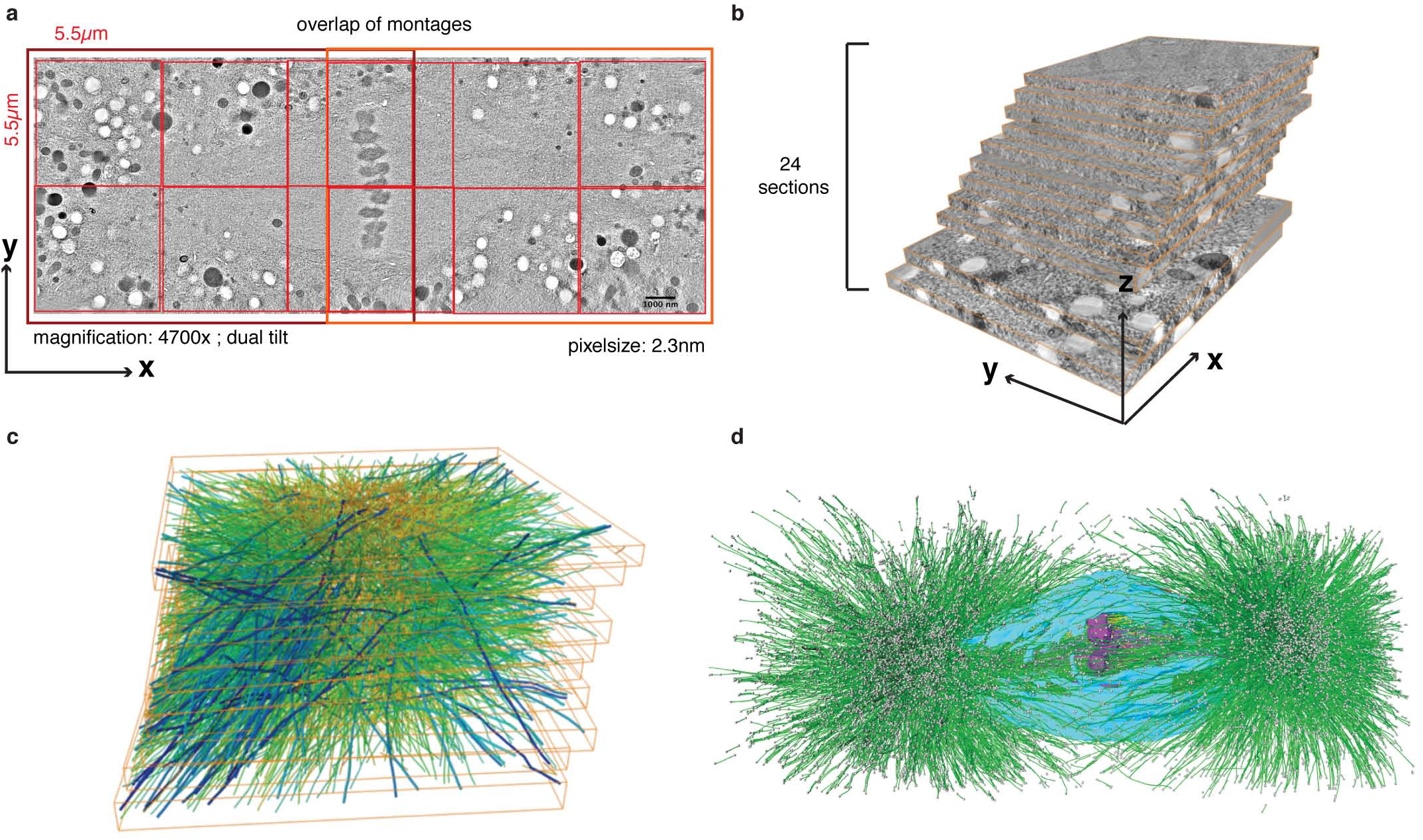
**Workflow of large-scale spindle reconstructions by 3D electron tomography** **a**, Two 2 × 3 montages (outlined in dark red, individual tomograms composing the montages are outlined in light red) are acquired and joined in X and Y to cover the entire area of the spindle. The size of a single tomogram, the magnification, and voxelsize are indicated. The thickness of a section is 300 nm **b**, Approximately 25 consecutive sections have to be acquired to cover the spindle volume. **c**, Microtubules (green) are automatically traced and manually corrected using the AMIRA software. This software is also used to stitch the individual sections in *z.* **d**, Features like chromosomes (purple) or the nuclear envelope (light blue) are segmented manually.

**Supplementary Figure 2.**
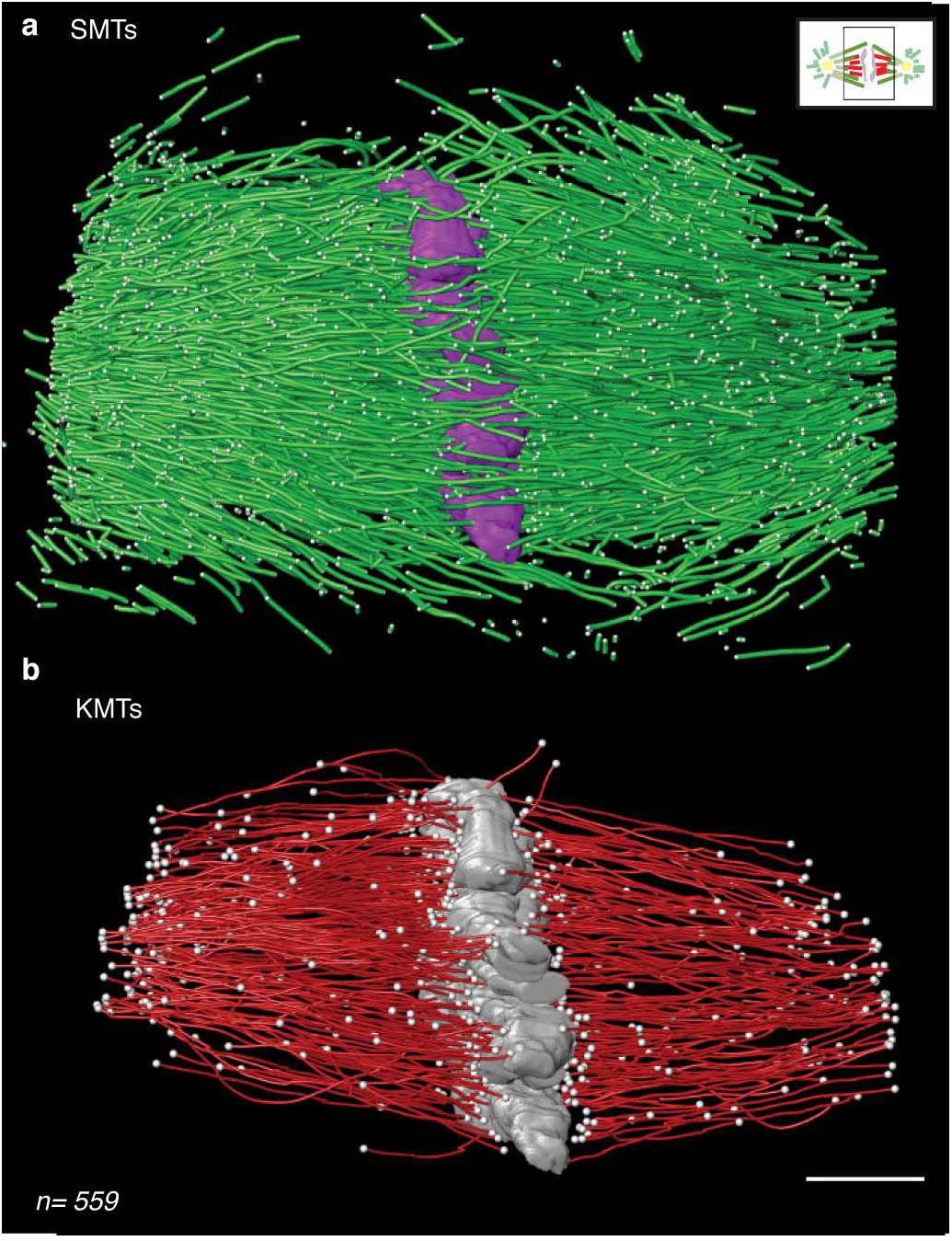
**Reconstruction of a central spindle in metaphase** **a**, Model of a metaphase spindle (Metaphase 3) covering the volume around the chromosomes. The region of the tomogram is indicated in the upper right corner. **b**, KMTs of the dataset as shown in **a.** The number of KMTs is indicated in the bottom left corner. Scale bar, 1 *μ*m.

**Supplementary Figure 3.**
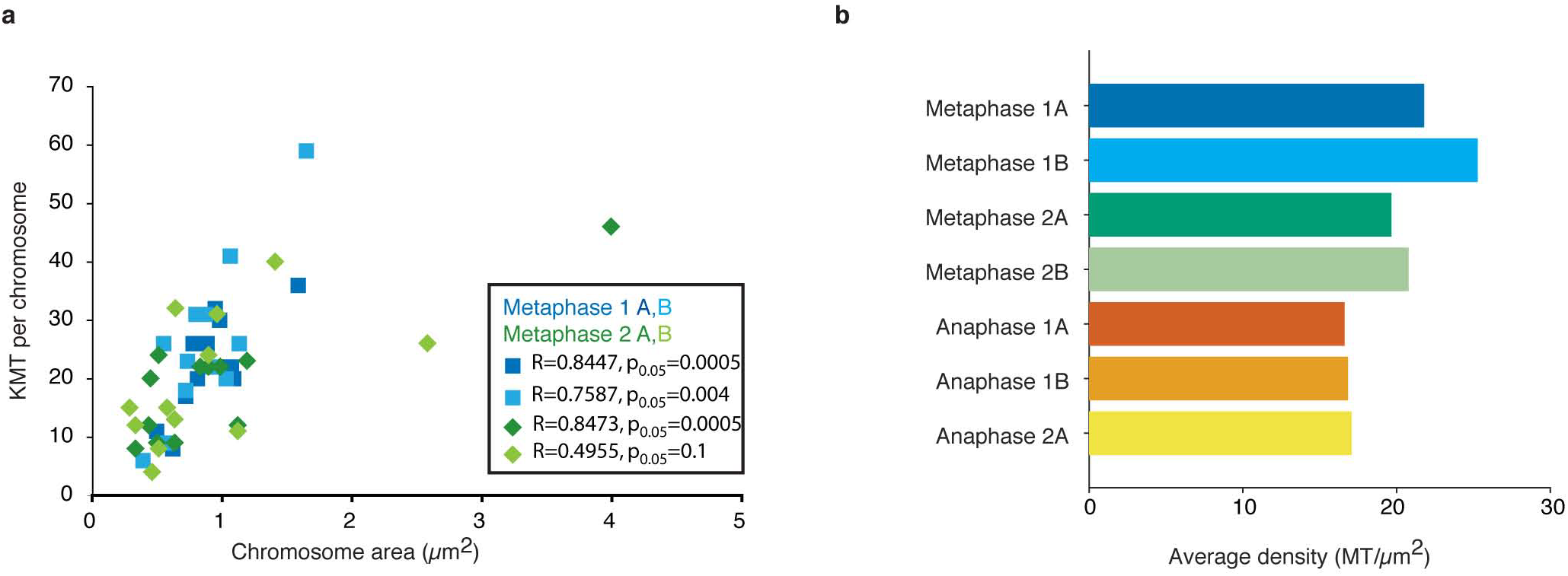
**KMT attachment correlates with chromosome area** **a**, Correlation of chromosome surface area and number of attached KMTs for two metaphase datasets. The Pearson’s correlation coefficient is indicated. **b**, Density of KMT attachment sites on chromosomes in metaphase and anaphase averaged over all chromosomes.

**Supplementary Figure 4.**
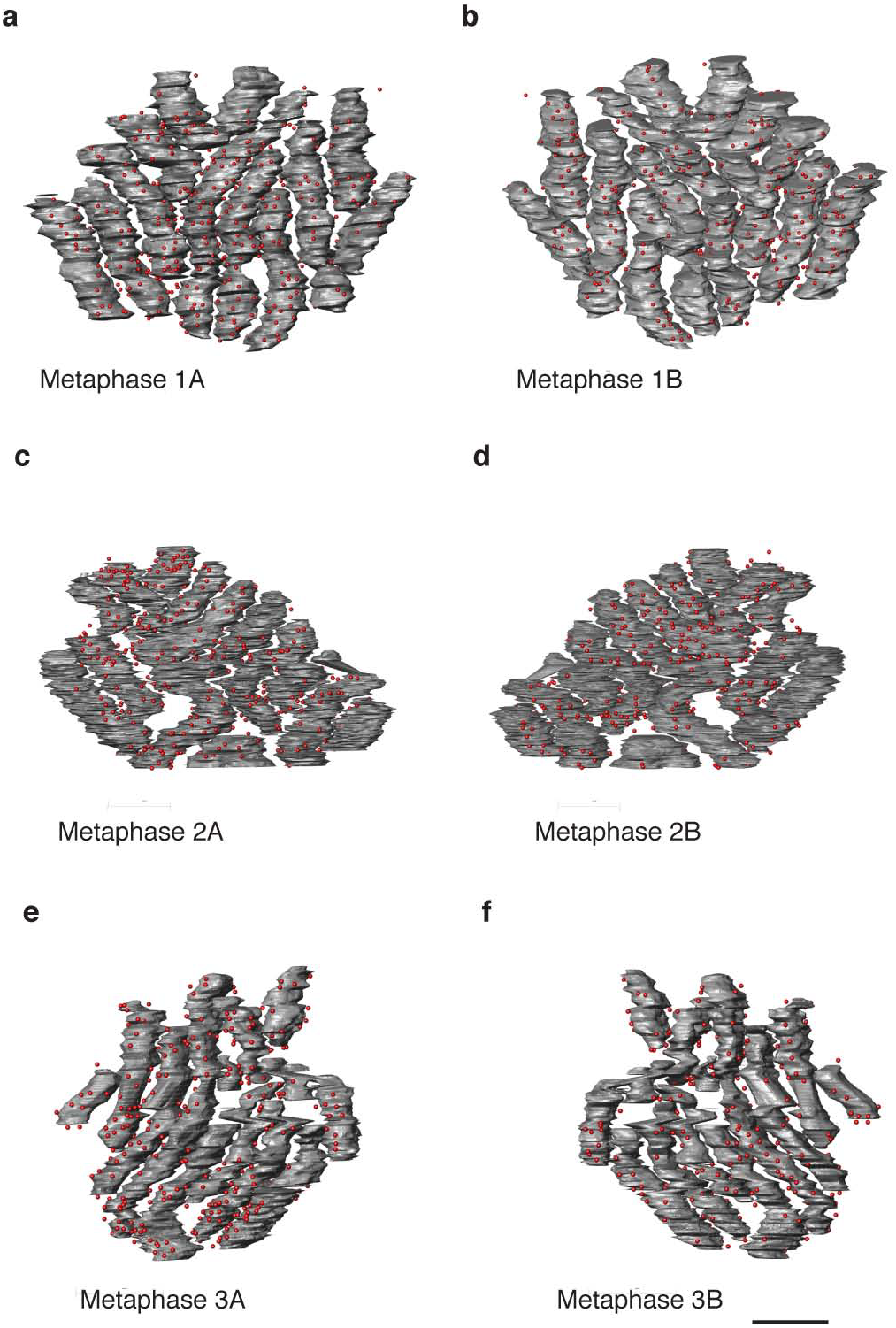
**KMT attachments sites on the chromosomes** End-on views of each metaphase plate as seen from both poles for the different datasets. **a, b**, Metaphase 1. **c, d**, Metaphase 2. **e, f**, Metaphase 3. Microtubule attachment to individual chromosomes from each pole is indicated by red dots. Scale bar, 1 *μ*m.

**Supplementary Figure 5.**
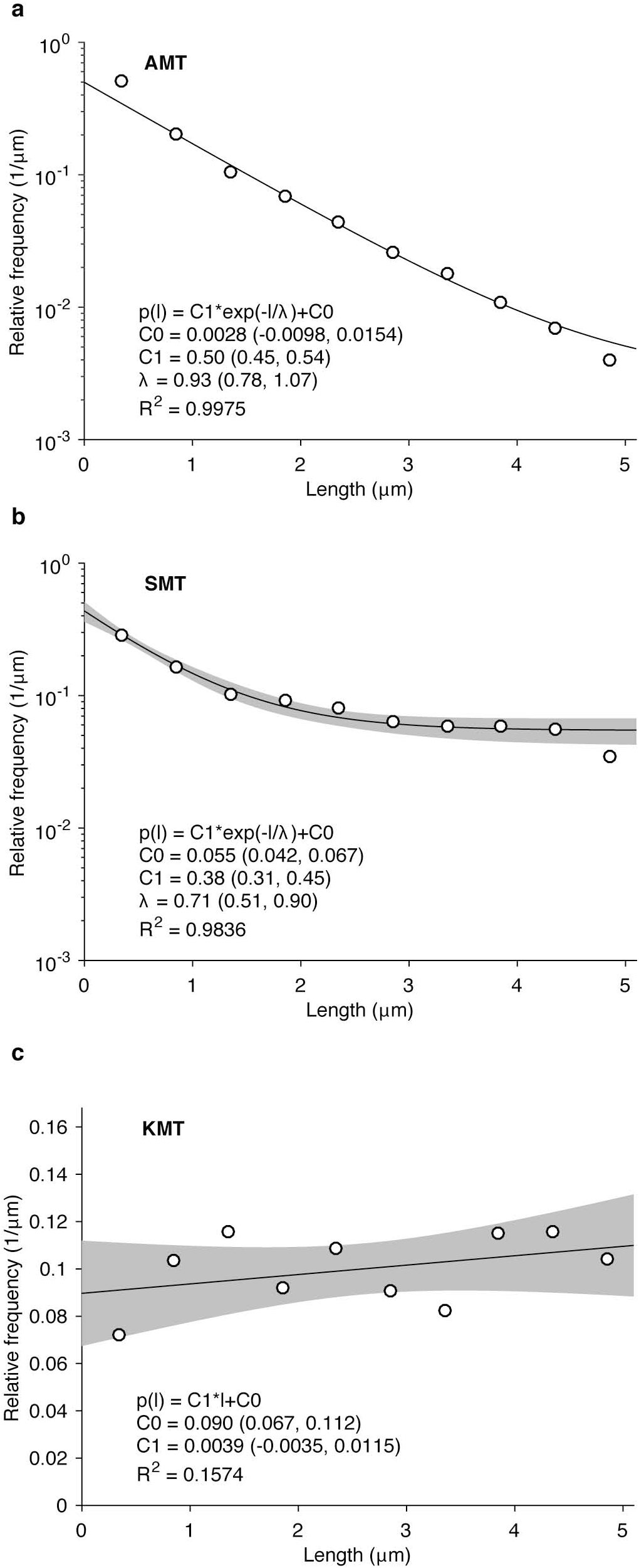
**Fitted histograms of the microtubule length distribution** **a**, Fit of the histogram of the AMT length distribution based on all five data sets to a single exponential with a constant. The gray shaded areas are the 95% confidence intervals for the fitted function for single observations. **b**, Fit of the histogram of the SMT length distribution based on all five data sets to a single exponential plus a constant. **c**, Fit of the histogram of the KMT length distribution based on all five data sets to a linear function. The fitting parameters are indicated with the 95% confidence intervals and the unadjusted coefficient of determination (R^2^) is provided.

**Supplementary Figure 6.**
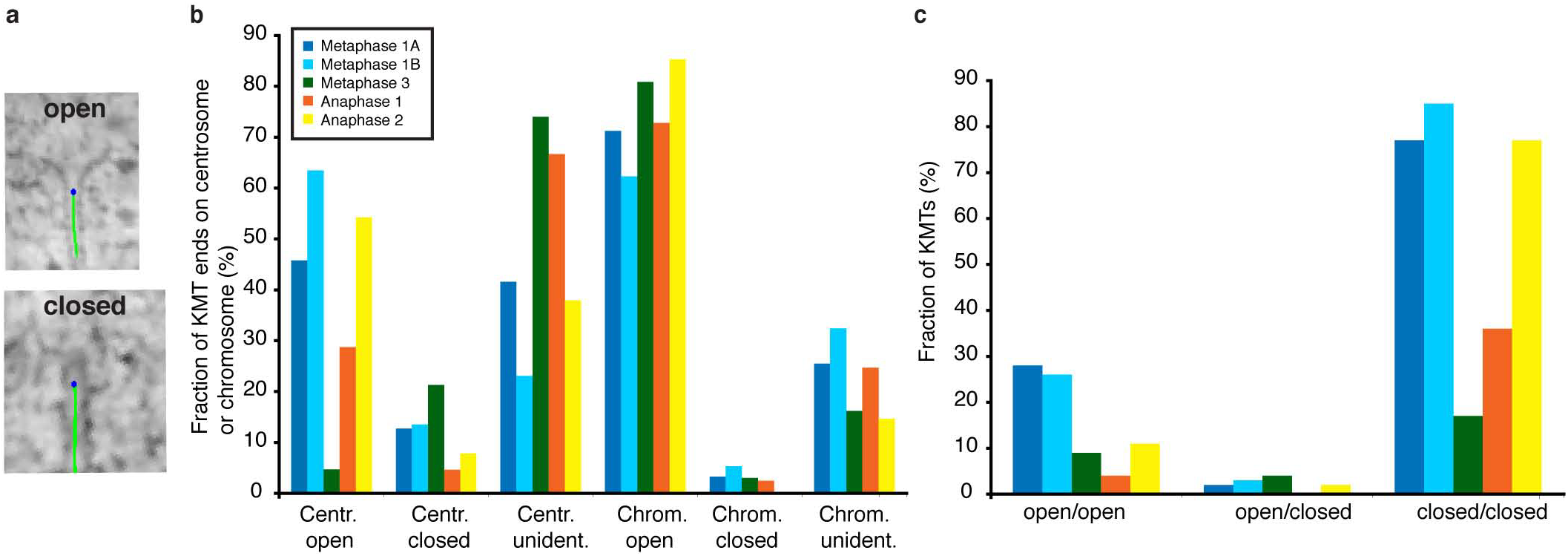
Analysis of microtubule end conformation **a**, Representative example for an open (upper panel) and closed microtubule end conformation (lower panel). **b**, Percentage of open, closed and unidentified KMT ends at the centrosomes and chromosomes in metaphase and anaphase. **c**, Percentage of conformations of both ends of individual KMTs in metaphase and anaphase.

**Supplementary Figure 7.**
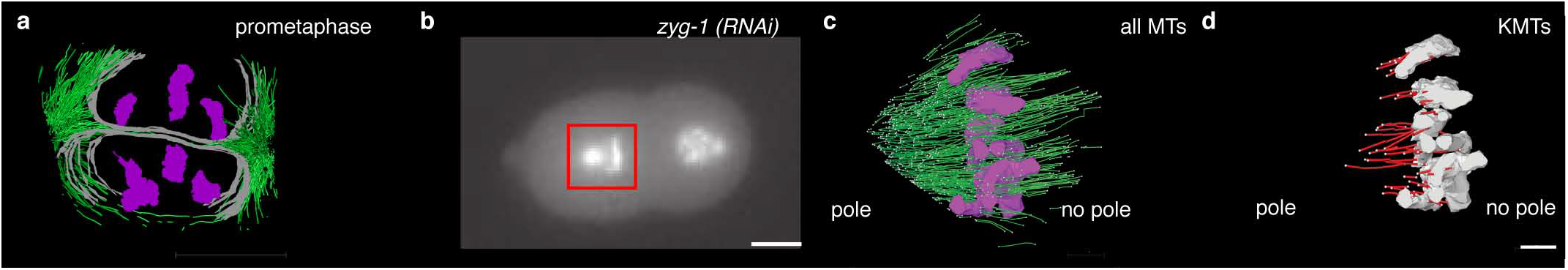
**Microtubules in early prometaphase and monopolar spindles** **a**, Model of chromosomes (magenta), nuclear envelope (grey) and microtubules (green) in a one-cell *C. elegans* embryo at early prometaphase. **b**, Two-cell *C. elegans* embryo after *zyg-1* (RNAi) labeled with β-tubulin::GFP and Histone::GFP. Red box indicates the area of the tomogram Scale bar, 10 *μ*m. **c**, Model of SMTs in three consecutive tomographic sections of a monopolar spindle as shown in **b. d**, Model of the KMTs as identified in **c.** Scale bar, 1 *μ*m.

**Supplementary Figure 8.**
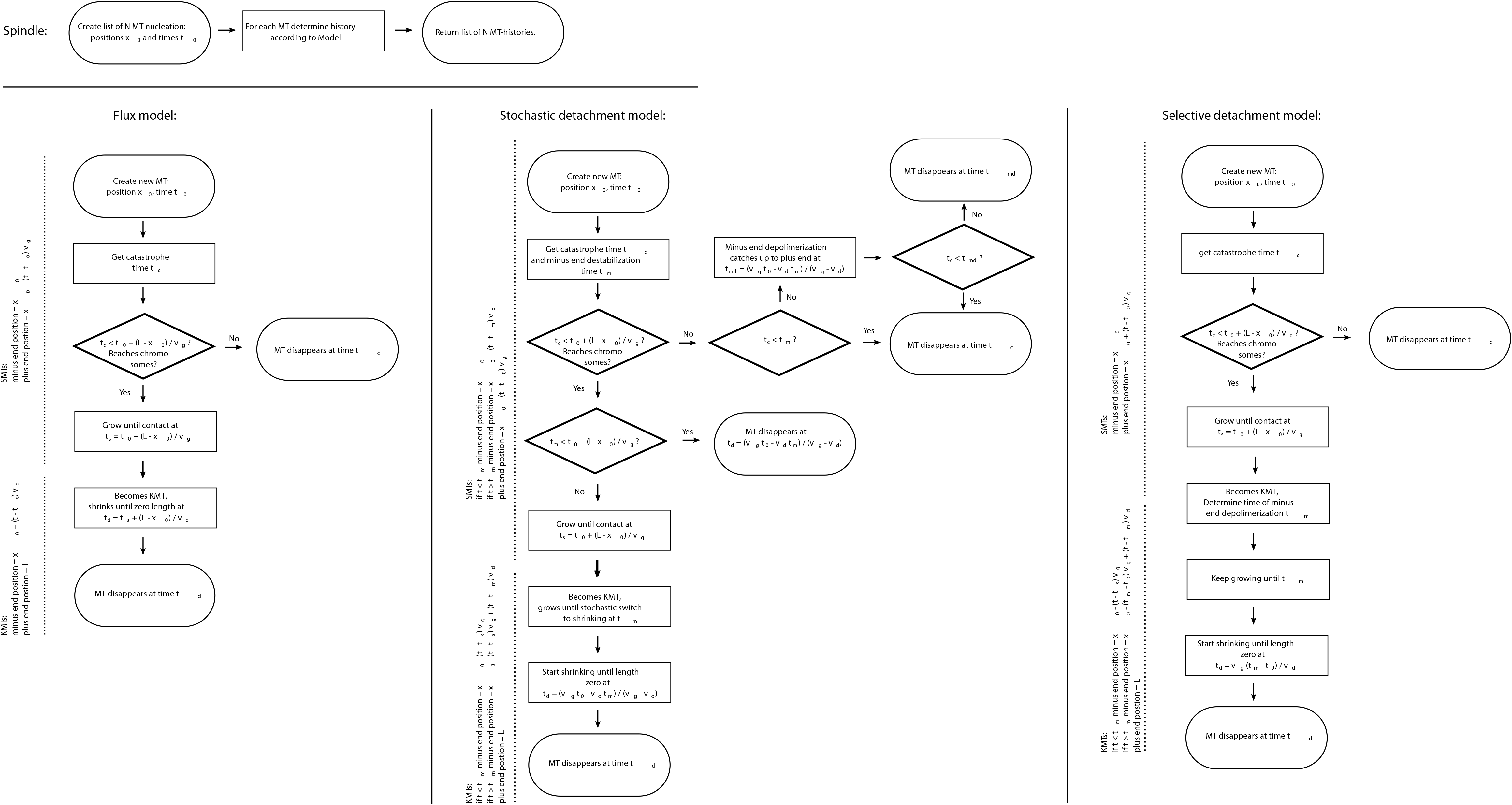
**Flowchart of the simulation process** Schematic flowchart depicting the different models in the context of a Gillespie algorithm.

**Supplementary Figure 9.**
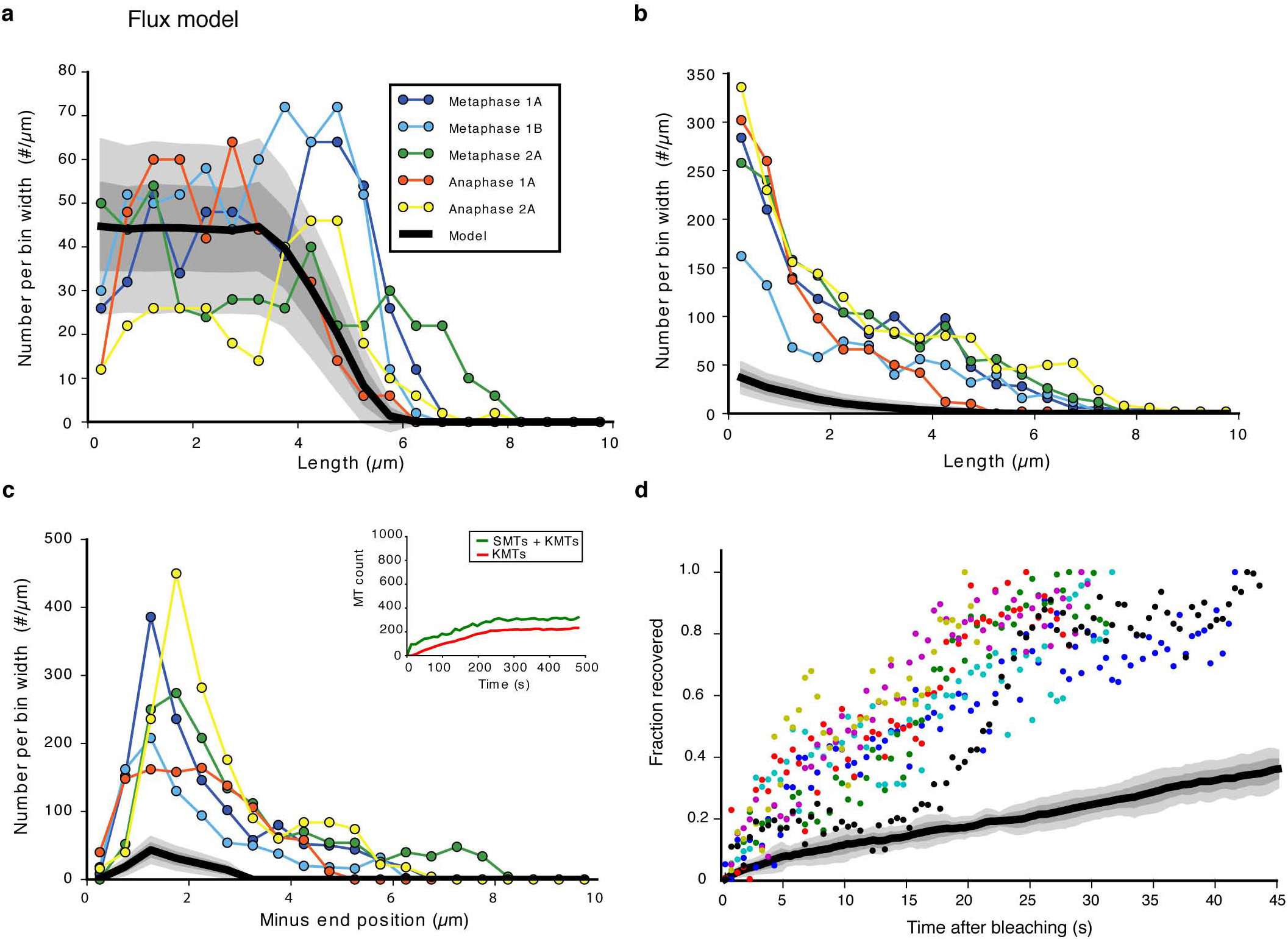
**Results from the stochastic microtubule *flux model*** **a**, KMT length distribution. **b**, SMT length distribution. **c**, SMT minus-end distribution. Inset shows the time-course of the total microtubule number (green) and KMT number (red) for a typical instance of the simulation with a depolymerisation velocity of V_d_ = 0.02 *μ*m/s **d**, Comparison of experimental FRAP data on microtubule recovery (individual measurement are shown in different colours) with the simulated FRAP data based on the *flux model.* For **a, b, c, d** we display the long expectation value of the model (solid black line) plus one (dark grey shaded region) and two (light grey shaded region) standard deviations.

**Supplementary Figure 10.**
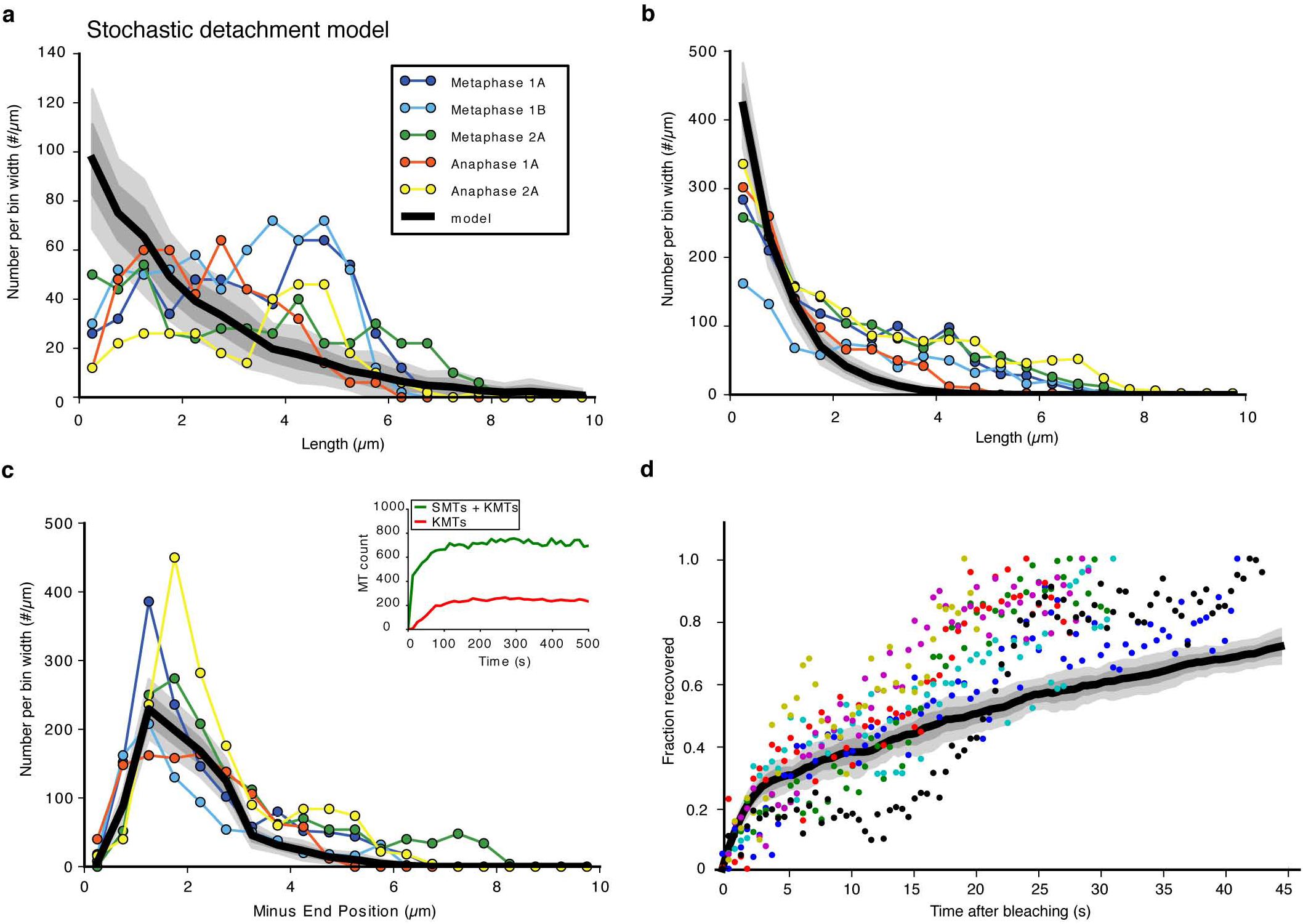
**Results from the *stochastic microtubule detachment model*** **a**, KMT length distribution. **b**, SMT length distribution. **c**, SMT minus end distribution. Inset shows the time-course of the total microtubule number (green) and KMT number (red) for a typical instance of the simulation with a depolymerisation velocity of V_d_ = 0.45 *μ*m/s and a switching rate from growth to shrinkage of r = 0.1 Hz. **d**, Comparison of experimental FRAP data on microtubule recovery (individual measurement are shown in different colours) with the simulated FRAP data based on the *stochastic detachment model.* For **a, b, c, d** we display the long expectation value of the model (solid black line) plus one (dark grey shaded region) and two (light grey shaded region) standard deviations.

**Supplementary Figure 11.**
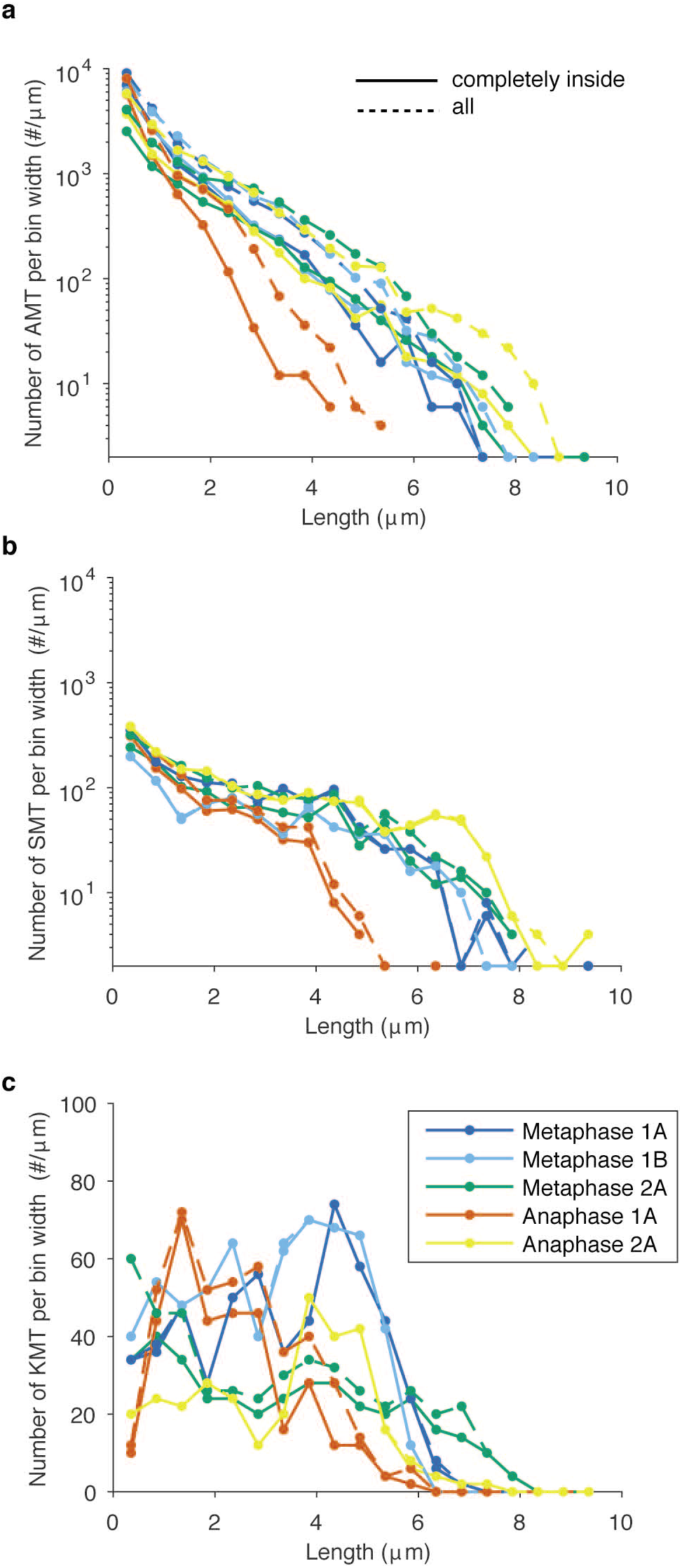
**Effect of the tomographic boundary on microtubule length distributions** **a**, Length distributions of all AMTs which are not touching the tomographic boundary and have both endpoints within the tomogram (solid line) and of all AMTs (dashed line). **b**, length distributions of all SMTs which are not touching the tomographic boundary and have both endpoints within the tomogram (solid line) and of all SMTs (dashed line). **c**, length distributions of all KMTs which are not touching the tomographic boundary and have both endpoints within the tomogram (solid line) and of all KMTs (dashed line).

**Supplementary Figure 12.**
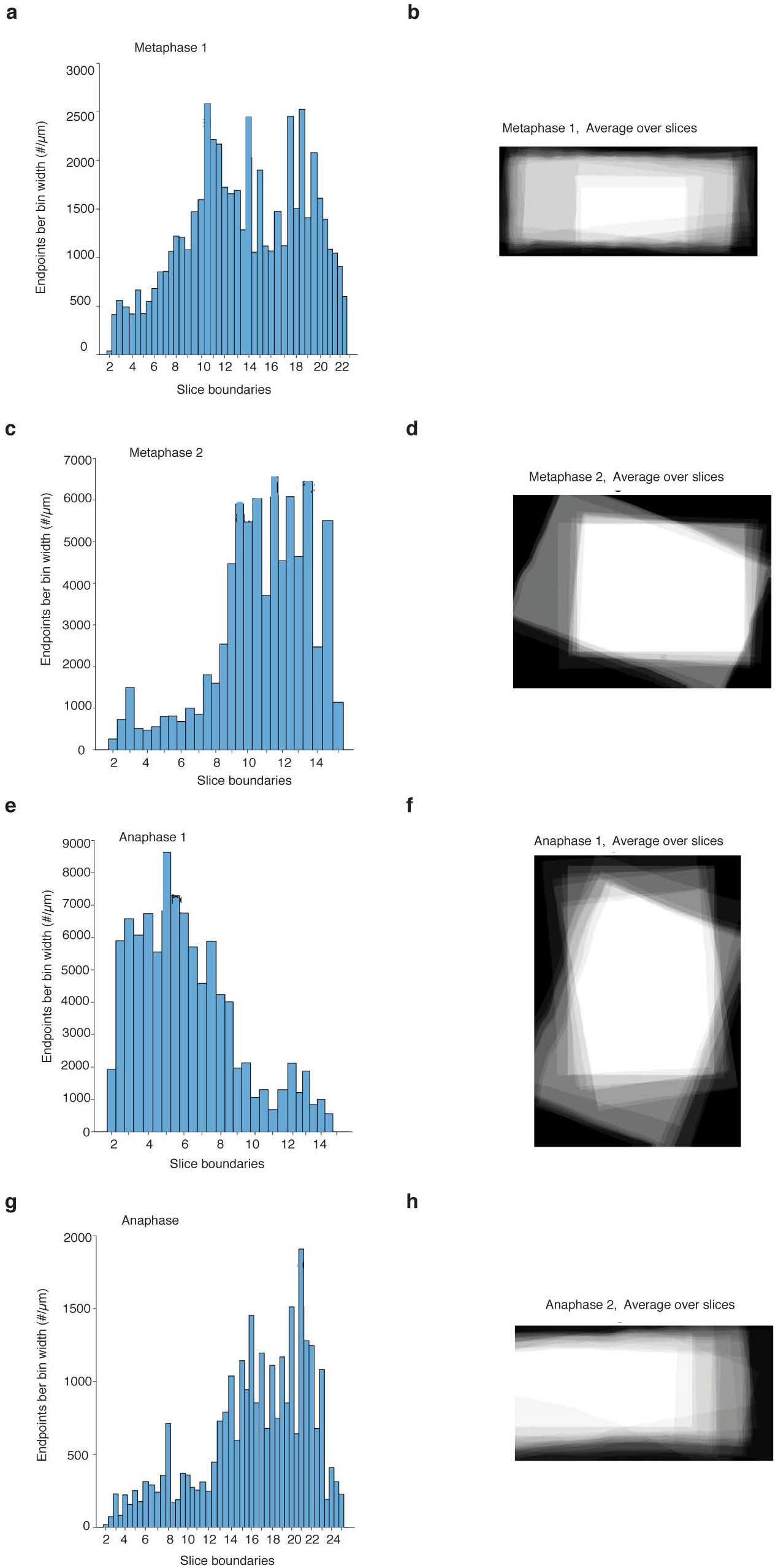
**Z-stack histograms of microtubule endpoints** **a,c,e,g**, Histograms of the number of microtubule endpoints in a volume, which is completely within a section and one across the boundary with comparable size. The number of endpoints is an estimate of the stitching quality of the individual datasets. Only endpoints in the intersection of all slices are analysed. **b,d,f,h**, Average of the tomogram area over all slices for each individual data set as shown in **a,c,e,g** as a z-projection.

**Supplementary Video 1. Visualization of the 3D reconstruction of a complete metaphase spindle in the early *C. elegans* embryo**

This full reconstruction (corresponding to Figure 1a) shows KMTs in red, and both AMTs and SMTs in green. The segmentation of the chromosomes is shown in blue.

**Supplementary Video 2. Close up-view of the chromosome region of a metaphase spindle in the early *C. elegans* embryo**

This movie (corresponding to Figure 1a) shows a close up-view of the microtubules around the metaphase plate. The rotation around the spindle axis shows KMTs in red, and both AMTs and SMTs in green. Chromosomes are visualized in blue.

**Supplementary Video 3. Visualization of microtubule plus-end growth in the metaphase spindle by EBP-2::GFP**

This movie (corresponding to Figure 4a,b) shows two examples of the motion of EBP-2::GFP comets in metaphase of the early *C.elegans* embryo. The exposure is 150 ms, the frame rate is 5 frames per second.

**Supplementary Video 4. Fluorescence recovery after photobleaching (FRAP) in *C. elegans* metaphase**

The FRAP experiment (corresponding to Figure 4c) in a histone::GFP and β-tubulin::GFP tagged *C. elegans* embryo in metaphase shows the recovery of the bleachmark over time. The exposure time is 100 ms, the frame rate is 2 frames per second.

**Supplementary Video 5. Laser microsurgery in the *C. elegans* metaphase spindle**

Laser microsurgery in a β-tubulin::GFP tagged *C. elegans* embryos was applied to induce the formation of novel microtubule plus and minus-ends. A single wave of microtubule depolymerisation can be observed. The exposure time is 300 ms and the frame rate is 1 frame every 5 seconds.

